# Structural basis of meiotic chromosome synaptic elongation through hierarchical fibrous assembly of SYCE2-TEX12

**DOI:** 10.1101/2020.12.30.424799

**Authors:** James M. Dunce, Lucy J. Salmon, Owen R. Davies

**Affiliations:** Institute of Cell Biology, University of Edinburgh, Michael Swann Building, Max Born Crescent, Edinburgh EH9 3BF; Biosciences Institute, Faculty of Medical Sciences, Newcastle University, Framlington Place, Newcastle upon Tyne NE2 4HH, UK; Department of Biochemistry, University of Cambridge, 80 Tennis Court Road, Old Addenbrookes Site, Cambridge CB2 1GA, UK

**Author notes:** **Author for Correspondence**, Owen Davies.

**Keywords:** Meiosis, chromosome structure, double-strand break, chiasmata, synaptonemal complex, SYCE2, TEX12, self-assembly, coiled-coil, small-angle X-ray scattering, biophysics, X-ray crystallography, intermediate filaments

## Abstract

The synaptonemal complex (SC) is a supramolecular protein assembly that mediates synapsis between homologous chromosomes during meiosis. SC elongation along the chromosome length (up to 24 μm) depends on its midline α-fibrous component SYCE2-TEX12. Here, we report X-ray crystal structures of SYCE2-TEX12 as an individual building-block and upon assembly within a fibrous lattice. We combine these structures with mutagenesis, biophysics and electron microscopy to reveal the hierarchical mechanism of SYCE2-TEX12 fibre assembly. SYCE2-TEX12’s building-blocks are 2:2 coiled-coils which dimerise into 4:4 hetero-oligomers and interact end-to-end and laterally to form 10-nm fibres, which intertwine within 40-nm bundled micrometre-long fibres that define the SC’s midline structure. This assembly mechanism bears striking resemblance with intermediate filament proteins vimentin, lamin and keratin. Thus, SYCE2-TEX12 exhibits behaviour typical of cytoskeletal proteins to provide an α-fibrous SC backbone that structurally underpins synaptic elongation along meiotic chromosomes.

## Introduction

Meiosis, the specialised form of cell division in gametogenesis, involves an extraordinary chromosome choreography that results in the formation of haploid oocytes and spermatozoa^1,2^. The hallmark of meiosis is synapsis, in which aligned homologous chromosomes become tightly bound along their axial length whilst undergoing recombination and crossover formation^2^. This unique state is achieved by the synaptonemal complex (SC), a supramolecular protein assembly that acts as a molecular ‘zipper’ of meiotic chromosomes^2,3^. Homologous chromosome synapsis by the SC is essential for generating healthy gametes by meiosis^4^, and mutations that disrupt SC structure have been identified in clinical cases of infertility^5,6^. Thus, the mechanism of SC assembly is fundamentally important for understanding human fertility and the molecular causes of infertility, miscarriage and aneuploidy.

In advance of synapsis, homologous chromosome pairs are established and aligned by a series of interhomologue recombination intermediates, which are generated by recombination searches from 200-400 double-strand breaks per cell^1,2,7^. Concomitant with recombination, chromosomes adopt a ‘lampbrush’ structure, in which chromatin is looped around linear axes^8–11^. Thus, the SC assembles between aligned linear arrays of homologous chromatin and converts discrete physical links of recombination into a single continuous synapsis along the axial length. Once assembled, the SC’s three-dimensional structure facilitates the resolution of recombination intermediates, with the formation of typically one crossover per arm^4,12^. In addition to providing genetic diversity, crossovers have a direct structural role in mediating the sole physical connections between homologues following SC disassembly, thereby enabling faithful homologous chromosome segregation at the end of the first meiotic division^1–3^.

The SC was discovered in 1956 through its iconic tripartite appearance on electron micrographs, initially in crayfish spermatocytes^13^, and subsequently throughout mammals, insects, plants and yeast^14,15^. The SC’s tripartite appearance of three parallel electron-dense fibres corresponds to the two chromosome axes (lateral elements), at a fixed 100-nm separation, with a midline central element (Figure 1a)^14,15^. The central and lateral elements have widths of approximately 20-40 nm and 50 nm, respectively, are continuous along the chromosome length, which is 4-24 μm in humans, and are held together by a series of interdigitated transverse filaments that act as the teeth of the SC ‘zipper‘^14–18^. Electron microscopy and super-resolution fluorescence microscopy have shown that the mammalian SC has a depth of up to 100 nm^17,19,20^. If we assume a square cross-section of 100-nm sides, with a solvent content between 20-80%, we can estimate that the combined supramolecular structure of the SC has a molecular weight of between 1.6-6.4 GDa per μm length. Thus, entire SCs may have molecular weights of between 6.4-154 GDa, placing them amongst the largest protein structures within a cell.

**Figure 1.**
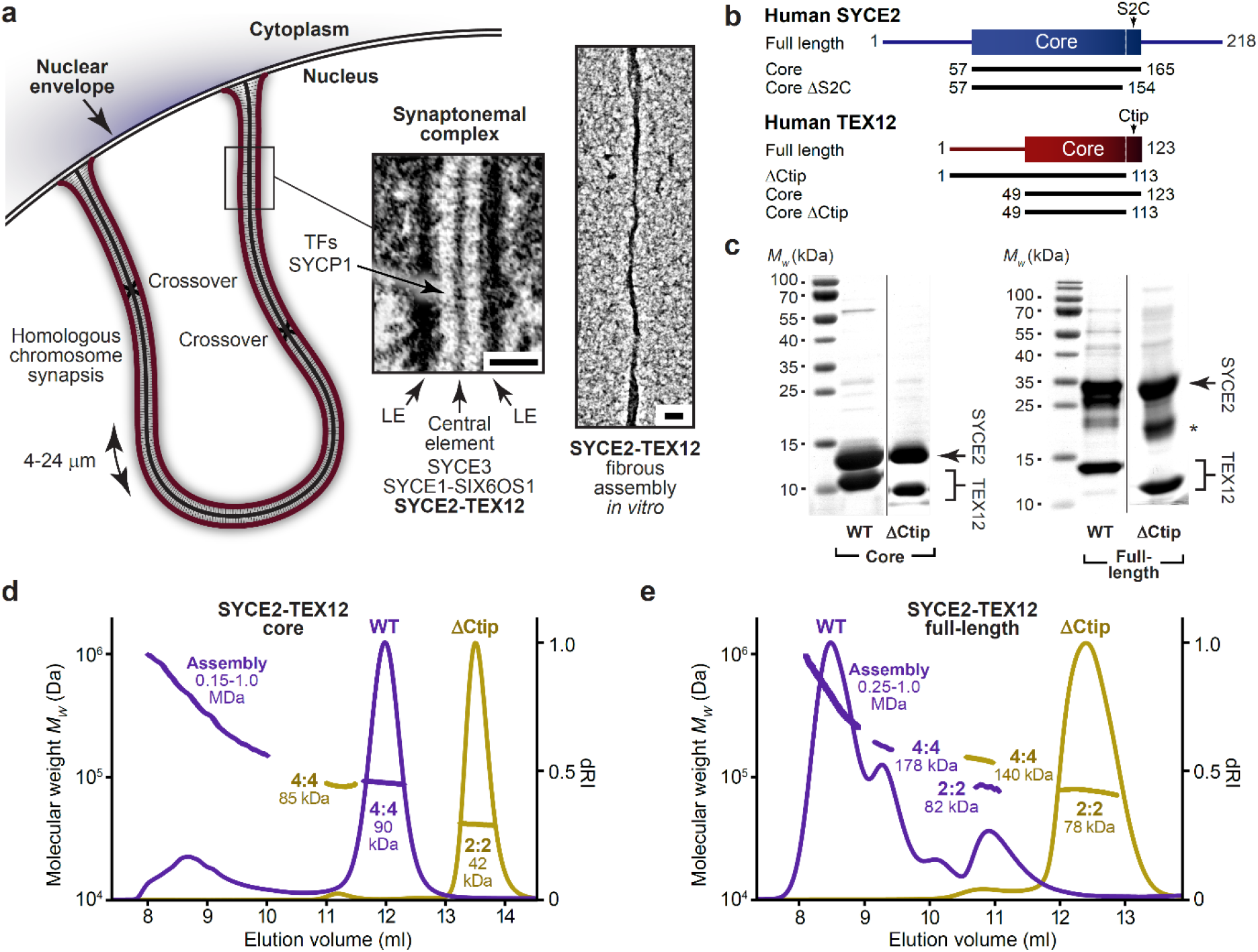
SYCE2-TEX12 self-assembly is driven by the C-terminal tip of TEX12. (**a**) Schematic of the synaptonemal complex (SC) mediating full synapsis between homologous chromosomes that are physically tethered at both ends to the nuclear envelope. The SC has a tripartite structure of an electron-dense central element lying midway between chromosome-bound lateral elements (electron micrograph reproduced from Kouznetsova, et al.^4^). SC central element formation depends on an SYCE2-TEX12 complex that undergoes fibrous self-assembly *in vitro* (electron micrograph reproduced from Davies, et al.^37^). Scale bars, 100 nm. (**b**) Human SYCE2 and TEX12 sequences, depicting their α-helical structural cores and the principal constructs used in this study. (**c**) SDS-PAGE of purified SYCE2-TEX12 full-length and core complexes containing wild-type (WT) and ΔCtip TEX12 sequences. The dominant degradation product of SYCE2 is indicated by an asterisk. (**d,e**) SEC-MALS analysis of SYCE2-TEX12 (**d**) core and (**e**) full-length complexes in which differential refractive index (dRI) is shown with fitted molecular weights (*M_w_*) plotted across elution peaks. (**d**) SYCE2-TEX12 core forms a 90 kDa 4:4 complex (75% of total mass; theoretical *Mw* – 89 kDa) and large molecular assemblies of up to 1.0 MDa (25%). Truncation of TEX12’s C-terminal tip (ΔCtip) restricts SYCE2-TEX12 assembly to a 42 kDa 2:2 complex (98%; theoretical – 42 kDa) and 85 kDa 4:4 complex (2%; theoretical – 84 kDa). (**e**) SYCE2-TEX12 full-length forms an 82 kDa 2:2 complex (12%; theoretical – 78 kDa), 178 kDa 4:4 complex (18%; theoretical – 156 kDa) and large molecular assemblies of up to 1.0 MDa (70%). The TEX12 ΔCtip truncation restricts SYCE2-TEX12 assembly to a 78 kDa 2:2 complex (98%; theoretical – 76 kDa) and 140 kDa 4:4 complex (2%; theoretical – 151 kDa).

Over the last thirty years, protein components of the mammalian SC have been identified and their roles defined by genetics and cellular imaging. Transverse filaments are formed by SYCP1 molecules, which are bioriented with their N- and C-termini within central and lateral elements, respectively, such that opposing molecules span the 100-nm width of the SC^17,20–23^. Lateral elements contain SYCP2 and SYCP3^24–26^, whilst the central element is formed of SYCE1, SYCE2, SYCE3, TEX12 and SIX6OS1^27–30^. SYCP1 and central element proteins are essential for SC formation and meiosis, with their individual mouse knockouts leading to infertility with failure of synapsis and crossover formation^28,30–34^. However, there are important subtle differences between these phenotypes. The SYCP1 knockout exhibits complete synaptic failure with no central element recruitment^28,30,31,34^. In SYCE1, SYCE3 and SIX6OS1 knockouts, there is patchy SYCP1 recruitment and some chromosomal associations, but without a tripartite SC^28–30,32^. in contrast, SYCE2 and TEX12 knockouts demonstrate short stretches of tripartite structure, containing SYCP1, SYCE1, SYCE3 and SIX6OS1, which fail to extend along the chromosome axis^33,34^. Thus, it has been proposed that SYCE1, SYCE3 and SIX6OS1 are initiation factors that provide shortrange stabilisation of nascent SYCP1 synapsis, whereas SYCE2 and TEX12 are elongation factors that underpin long-range SC growth by providing longitudinal structural support^28,33–35^. This dichotomy is underpinned by biochemical findings that SYCE1 directly interacts with SYCE3 and SIX6OS^15,36^, whilst SYCE2 and TEX12 exist in a seemingly constitutive complex^37^.

In the last decade, the molecular underpinnings of the SC have started to come into focus through analysis of SC proteins by structural biology^5,23,36–43^. An emerging theme is that SC proteins, which are almost universally coiled-coils, exist as defined building-blocks that self-assemble into higher-order structures to provide the SC’s distinct architectural elements. SYCP1 is a tetramer that assembles into a lattice-like array to fulfil the fundamental role of the SC in tethering chromosome axes^23^. SYCP3 is a tetramer that assembles into paracrystalline fibres, which have been observed for the recombinant protein *in vitro*^38,43^, upon heterologous cellular expression^44–46^, and in the native mammalian SC^47^. SYCP3 fibres separate DNA-binding sites by a 23-nm repeating unit along the longitudinal axis, which facilitates the compaction of chromatin loops within the meiotic chromosome axis^38^. Central element proteins SYCE2 and TEX12 form a core 4:4 complex that assembles into fibres with polymorphic appearance, of 40-nm width, and up to 5-μm in length (Figure 1a)^37^. The appearance and dimensions of SYCE2-TEX12 fibres match those of the native SC central element, suggesting that fibrous assembly along the SC axis underlies the structural role of SYCE2-TEX12 in synaptic elongation.

Here, we combine X-ray crystal structures of SYCE2-TEX12 at two critical stages of assembly, with solution and electron microscopy studies, to reveal a hierarchical mechanism for SYCE2-TEX12 α-fibre assembly, which intriguingly resembles intermediate filament assembly. The building-blocks of SYCE2-TEX12 assembly are 2:2 complexes, which dimerise into 4:4 hetero-octamers that undergo end-to-end and staggered lateral interactions to form 10-nm fibres. These thread-like structures become intertwined within 40-nm fibres that extend to several micrometres in length and resemble the native SC central element. Thus, we define the molecular mechanism whereby SYCE2-TEX12 forms an α-fibrous axis that provides long-range structural support for synapsis as the ‘backbone’ of the SC.

## Results

### SYCE2-TEX12 fibre assembly is mediated by TEX12’s C-terminal tip

A common feature of self-assembly by SC proteins SYCP1 and SYCP3 is the presence of short motifs at the ends of their α-helical cores that direct higher-order assembly^23,38^. In each case, disruption of these motifs blocks assembly and restricts proteins to their obligate ‘building-block’ structures. Could a similar phenomenon apply to SYCE2-TEX12 fibres? To test this, we analysed the oligomeric state and assembly of short deletions of the SYCE2-TEX12 core (amino-acids 57-165 and 49-123; Figure 1b) by size-exclusion chromatography multi-angle light scattering (SEC-MALS). Having confirmed that wildtype SYCE2-TEX12 core is a 4:4 complex that assembles into higher molecular weight species (Figure 1c,d), we identified that deletion of ten amino-acids at TEX12’s C-terminal tip (ΔCtip, amino-acids 49-113; Figure 1b) restricted the SYCE2-TEX12 complex to a 2:2 stoichiometry and blocked its higher-order assembly (Figure 1c,d). Circular dichroism (CD) determined that wild-type and ΔCtip complexes have comparable α-helical contents (93% and 91%) and melting temperatures (71°C and 67°C) (Supplementary Figure 1a,b). Thus, the ΔCtip 2:2 complex may constitute a substructure of the wildtype core 4:4 complex, and hence may represent its minimum building-block for assembly.

We find that full-length SYCE2-TEX12 has an increased propensity for assembling in solution, with SEC-MALS revealing its predominant formation of higher-order species, in addition to clearly discernible 4:4 and 2:2 complexes (Figure 1c,e). The presence of a wild-type 2:2 complex supports its putative role as the minimum obligate SYCE2-TEX12 structure that undergoes hierarchical assembly into the observed 4:4 and higher-order species. In common with the core complex, deletion of TEX12’s C-terminal tip restricted full-length SYCE2-TEX12 to a 2:2 complex and blocked higher-order assembly (Figure 1c,e). The α-helical structure and stability of the wild-type protein was retained in the ΔCtip complex (Supplementary Figure 1a,c). Thus, in a manner reminiscent of SYCP1 and SYCP3^23,38^, deletion of a short motif at the C-terminal tip of TEX12 blocks higher-order assembly and stabilises SYCE2-TEX12 in its building-block 2:2 complex.

We next visualised SYCE2-TEX12 assembly by electron microscopy (Figure 2). This confirmed that core and full-length complexes assemble into morphologically similar fibres, which are typically positively-stained, vary in thickness up to approximately 40 nm and extend up to several micrometres in length (Figure 2a,d). Their distributions were determined through automated detection and measurement of fibres (Supplementary Figure 2), revealing that despite their similar lengths, mean fibre widths were slightly smaller for the full-length complex (12.8 ± 3.1 nm; Figure 2e,f) than its structural core (16.9 ± 4.1 nm; Figure 2b,c). This suggests that flanking sequences may provide additional interactions that confer stability to narrower core assemblies. In agreement with our SEC-MALS analysis, the TEX12 ΔCtip deletion blocked fibre formation of core and full-length complexes, as determined by manual inspection (Figure 2a,d) and automated detection (Figure 2c,f). We conclude that TEX12’s C-terminal tip is essential for assembly of SYCE2-TEX12 fibres, and its deletion stabilises the building-block 2:2 structure, providing a critical means for elucidating the fibre assembly mechanism.

**Figure 2.**
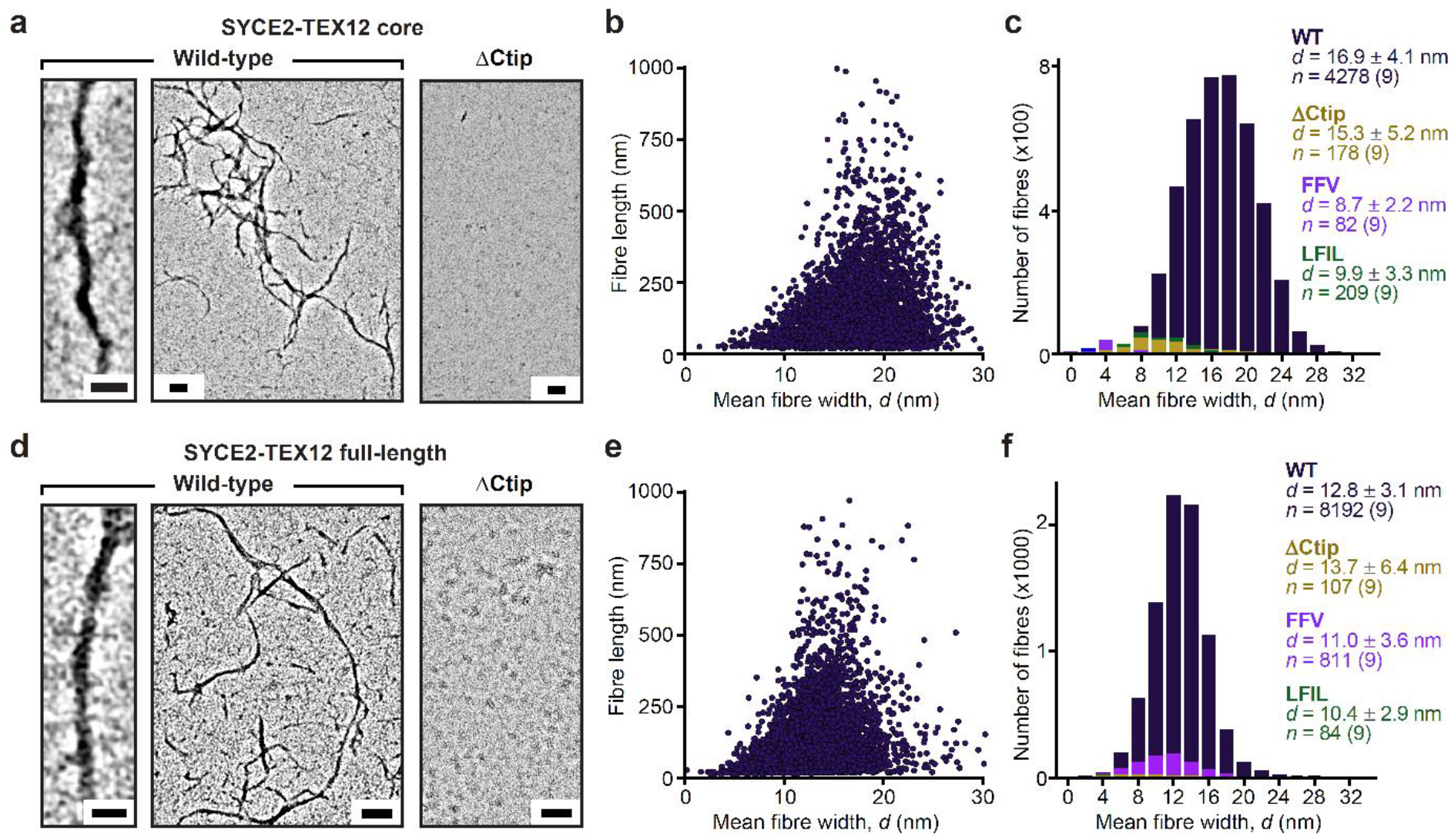
Fibrous assembly of SYCE2-TEX12. (**a-f**) Electron microscopy of SYCE2-TEX12 (**a-c**) core and (**d-f**) wild-type. (**a**) Electron micrographs of SYCE2-TEX12 core wild-type and ΔCtip. Scale bars, 50 nm (left) and 100 nm (middle and right). (**b**) Scatter plot of mean fibre width (*d*, nm) against fibre length (nm) for wild-type SYCE2-TEX12 core. (**c**) Histograms of the number of fibres with mean widths within 2-nm bins for SYCE2-TEX12 core wildtype (dark blue), ΔCtip (yellow), FFV (F102A, F109A, V116A; light blue) and LFIL (L110E, F114E, I117E, L121E; green). The mean, standard deviation and number of fibres in each population is shown, each determined from nine micrographs. (**d**) Electron micrographs of SYCE2-TEX12 full-length wild-type and ΔCtip. Scale bars, 25 nm (left) and 100 nm (middle and right). (**e**) Scatter plot of mean fibre width (d, nm) against fibre length (nm) for wild-type SYCE2-TEX12. (**f**) Histograms of the number of fibres with mean widths within 2-nm bins for SYCE2-TEX12 full-length wild-type (dark blue), ΔCtip (yellow), FFV (light blue) and LFIL (green). The mean, standard deviation and number of fibres in each population is shown, each determined from nine micrographs.

### Crystal structure of SYCE2-TEX12 in a 2:2 complex

The enhanced stability of the SYCE2-TEX12 ΔCtip core complex enabled the growth of protein crystals, which diffracted anisotropically to 2.42-3.36 Å resolution and enabled structure solution by molecular replacement of ideal helical fragments using *ARCIMBOLDO*^43^ (Table 1 and Supplementary Figure 3a). This revealed a 2:2 helical assembly in which an anti-parallel SYCE2 dimer spans the 14-nm length of the molecule and positions TEX12 chains in a staggered configuration (Figure 3a-c). The TEX12 chains are arranged with their N-termini towards the midline and their C-termini at either end of the molecule, immediately adjacent to the N-and C-termini of the SYCE2 antiparallel dimer (Figure 3a-c). The resultant 2:2 molecule has an unusual architecture in which a central four-helical bundle, formed of the SYCE2 dimer and overlapping N-terminal TEX12 chains, is flanked by three-helical bundles, each formed of the SYCE2 dimer and a single C-terminal TEX12 chain (Figure 3a,d-f). Importantly, the C-terminal ends of SYCE2’s core were not visible in electron density. Thus, deletion of TEX12’s C-terminal tip corresponded with disorder of an analogous C-terminal sequence of the SYCE2 core (herein referred to as S2C; amino acids 155-165), such that the molecular ends of the 2:2 complex are formed of closely associated truncated C-termini of SYCE2 and TEX12 (Figure 3a,c).

**Figure 3.**
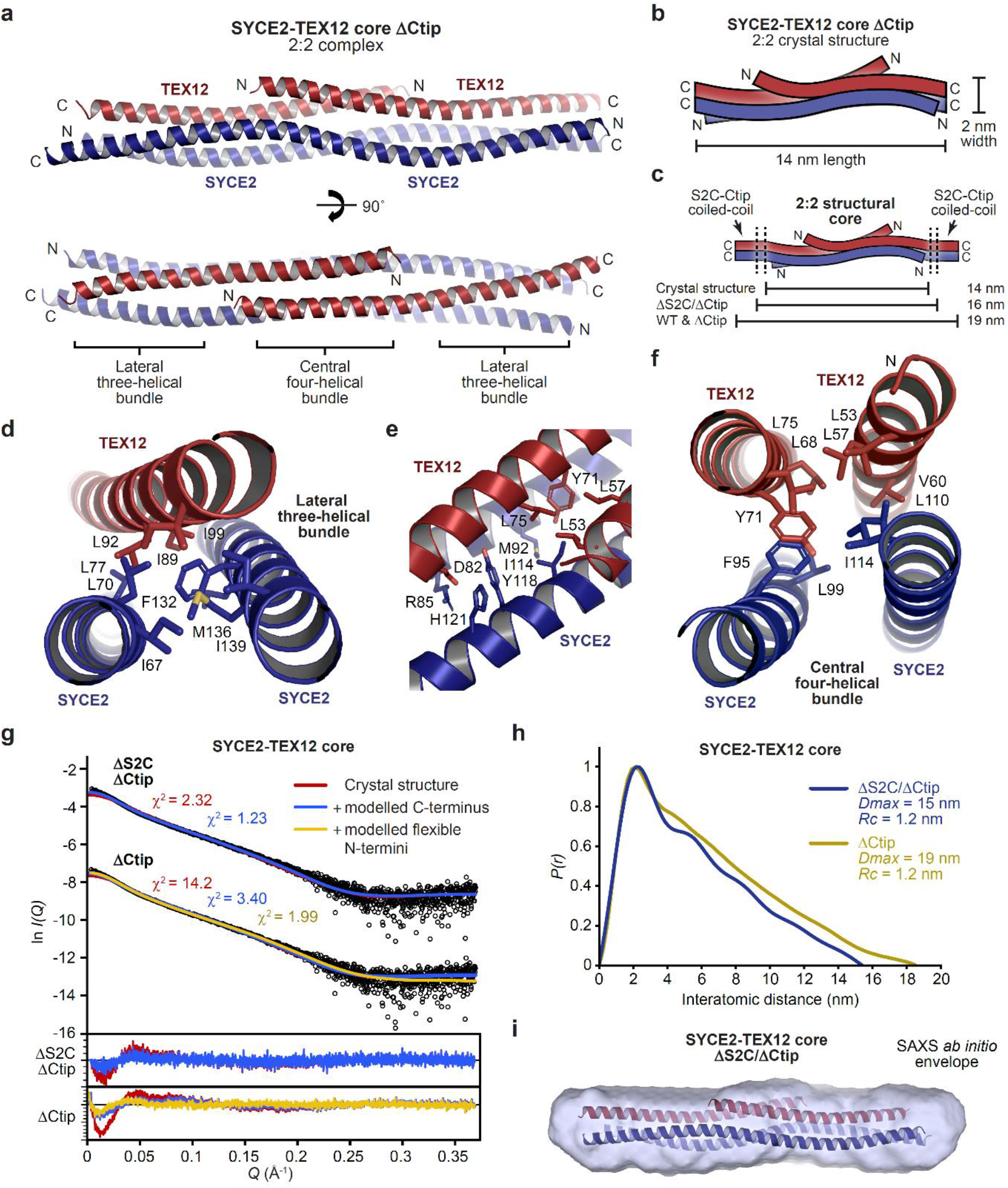
Crystal structure of the SYCE2-TEX12 core 2:2 complex. (**a**) Crystal structure of SYCE2-TEX12 core ΔCtip as a 2:2 complex. The two TEX12 chains (red) are bound to either end of an SYCE2 anti-parallel dimer (blue), with their C-termini oriented externally and immediately adjacent to SYCE2’s C-termini. The structure consists of a central four-helical bundle flanked by lateral three-helical bundles. (**b-c**) Schematics of the 2:2 crystal structure in which (**b**) the orientation of SYCE2 and TEX12 chains are highlighted, and (**c**) their additional S2C and Ctip sequences are shown as C-terminal coiled-coils. The 2:2 structure has a width of 2 nm and a length of 14 nm, which is predicted to increase to 19 nm upon addition of S2C and/or Ctip sequences. (**d**) The lateral three-helical bundle consists of an anti-parallel arrangement of SYCE2 chains and a single TEX12 chain, with a hydrophobic core formed by the residues indicated. (**e**) The transition point between lateral three-helical bundle and central four-helical bundle. (**f**) The central four-helical bundle consists of an anti-parallel arrangement of two TEX12 and two SYCE2 chains, with a hydrophobic core formed by the residues indicated. (**g-i**) SEC-SAXS analysis of SYCE2-TEX12 core ΔS2C/ΔCtip and ΔCtip. (**g**) SAXS scattering data overlaid with the theoretical scattering curves of the 2:2 crystal structure (red), with modelled C-terminal coiled-coil and S2C helix (blue), and with flexibly modelled N-termini (yellow); χ^2^ values are indicated and residuals for each fit are shown (inset). (**h**) SAXS *P(r)* interatomic distance distributions of SYCE2-TEX12 core ΔS2C/ΔCtip and ΔCtip, showing maximum dimensions *(Dmax)* of 15 nm and 19 nm, respectively. Their cross-sectional radii (*Rc*) were determined as 1.2 nm. (**i**) SAXS *ab initio* model of SYCE2-TEX12 core ΔS2C/ΔCtip. A filtered averaged model from 30 independent DAMMIF runs is shown with the 2:2 crystal structure docked into the SAXS envelope.

**Table 1.** Data collection, phasing and refinement statistics

| | SYCE2-TEX12 ΔCtip<br>2:2 complex | SYCE2-TEX12 ΔCtip<br>4:4 assembly |
| --- | --- | --- |
| **PDB accession** | 6R17 | 6YQF |
| **Data collection** |  |  |
| Space group | P2<sub>1</sub> | P2<sub>1</sub>2<sub>1</sub>2 |
| Cell dimensions |  |  |
| *a*, *b*, *c* (Å) | 88.52, 24.19, 88.48 | 42.67, 59.68, 156.49 |
| $\alpha$, $\beta$, $\gamma$ (°) | 90, 115.737, 90 | 90, 90, 90 |
| Resolution (Å) | 79.74 – 2.42 (2.74 – 2.42)* | 47.45 – 3.33 (3.59 – 3.33) |
| Ellipsoidal resolution (Å) | 2.416 (0.780 *a* * - 0.625 *c* *) | 4.203 (*a* *) |
| (direction) | 2.795 (*b* *) | 3.710 (*b* *) |
|  | 2.926 (0.491 *a* * + 0.871 *c* *) | 2.891 (*c* *) |
| *R*<sub>meas</sub> | 0.074 (1.293) | 0.246 (2.445) |
| *R*<sub>pim</sub> | 0.049 (0.914) | 0.098 (0.906) |
| *I* / $\sigma$(*I*) | 10.2 (1.2) | 4.7 (1.0) |
| *CC*<sub>1/2</sub> | 0.999 (0.564) | 0.995 (0.516) |
| Completeness (spherical) (%) | 63.2 (10.5) | 64.8 (27.1) |
| Completeness (ellipsoidal) (%) | 86.2 (51.3) | 83.0 (82.7) |
| Redundancy | 3.4 (3.0) | 6.5 (7.2) |
| **Refinement** |  |  |
| Resolution (Å) | 79.74 – 2.42 | 47.45 – 3.33 |
| No. reflections | 8497 | 4053 |
| *R*<sub>work</sub> / *R*<sub>free</sub> | 0.2871/0.3296 | 0.3266/0.3672 |
| No. atoms | 2536 | 2426 |
| Protein | 2526 | 2426 |
| Ligand/ion | 0 | 0 |
| Water | 10 | 0 |
| *B*-factors | 76.79 | 56.52 |
| Protein | 76.89 | 56.52 |
| Ligand/ion | N/A | N/A |
| Water | 53.44 | N/A |
| R.m.s. deviations |  |  |
| Bond lengths (Å) | 0.005 | 0.001 |
| Bond angles (°) | 0.788 | 0.272 |

\*Values in parentheses are for highest-resolution shell.

Does the 2:2 crystal structure represent the solution state of SYCE2-TEX12 ΔCtip? Size-exclusion chromatography small-angle X-ray scattering (SEC-SAXS) of the ΔCtip core 2:2 complex revealed a real-space *P(r)* interatomic distance distribution and Guinier analysis indicative of a rod-like molecule with 19-nm length and 1.2-nm cross-sectional radius (Figure 3g,h, Table 2 and Supplementary Figure 3b,c). These dimensions match the length of the crystal structure with missing S2C ends modelled as ideal helices, and its observed 2-nm width (Figure 3c and Supplementary Figure 3e). Further, the SAXS scattering curve was closely fitted by the crystal structure with modelled S2C helices and flexible N-termini (χ^2^ = 1.99; Figure 3g and Supplementary Figure 3e). We also analysed a ΔS2C/ΔCtip complex in which the C-termini of SYCE2 and TEX12 were deleted. SEC-MALS confirmed that the ΔS2C/ΔCtip core complex is predominantly 2:2 in solution (Supplementary Figure 3d). SEC-SAXS determined a rodlike molecule with 15-nm length and 1.2-nm cross-sectional radius, with an *ab initio* envelope that matches the dimensions of the 2:2 crystal structure (Figure 3g-i, Table 2 and Supplementary Figure 3c). Further, the SAXS scattering curve was closely fitted by the crystal structure upon ideal helical modelling of the few missing C-terminal amino-acids (χ^2^ = 1.23; Figure 3g). We conclude that the 2:2 crystal structure corresponds to the solution state of the ΔCtip core complex and hence represents the obligate 2:2 structure of SYCE2-TEX12 that acts as its building-block for assembly.

**Table 2.** Summary of SEC-SAXS data

| SYCE2-TEX12<br>core | WT<br>4:4 | FVF<br>4:4 | LFIL<br>2:2 | ΔCtip<br>2:2 | ΔS2C<br>ΔCtip<br>2:2 | WT<br>35 nm (I)<br>fibre | WT<br>65 nm (I)<br>fibre |
| --- | --- | --- | --- | --- | --- | --- | --- |
| **Guinier analysis** |  |  |  |  |  |  |  |
| $I(0)$ (cm<sup>-1</sup>) | 0.019<br>$\pm 1.4 \times 10^{-4}$ | 0.120<br>$\pm 2.0 \times 10^{-4}$ | 0.019<br>$\pm 4.5 \times 10^{-5}$ | 0.044<br>$\pm 1.3 \times 10^{-4}$ | 0.021<br>$\pm 5.0 \times 10^{-5}$ | 0.031<br>$\pm 7.4 \times 10^{-4}$ | 0.042<br>$\pm 6.2 \times 10^{-3}$ |
| $R_g$ (Å) | 49<br>$\pm 0.33$ | 49<br>$\pm 1.16$ | 47<br>$\pm 0.59$ | 49<br>$\pm 0.31$ | 41<br>$\pm 1.69$ | 86<br>$\pm 1.30$ | 167<br>$\pm 25.3$ |
| $R_c$ (Å) | 19.4 | 19.4 | 12.4 | 12.2 | 11.7 | 16.6 | 15.3 |
| $q_{min}$ (Å<sup>-1</sup>) | 0.056 | 0.0036 | 0.0070 | 0.0039 | 0.0043 | 0.0042 | 0.0035 |
| **$P(r)$ analysis** |  |  |  |  |  |  |  |
| $I(0)$ (cm<sup>-1</sup>) | 0.019<br>$\pm 3.2 \times 10^{-5}$ | 0.122<br>$\pm 2.0 \times 10^{-4}$ | 0.020<br>$\pm 3.9 \times 10^{-5}$ | 0.044<br>$\pm 1.1 \times 10^{-4}$ | 0.021<br>$\pm 6.2 \times 10^{-5}$ | 0.031<br>$\pm 6.3 \times 10^{-4}$ | 0.045<br>$\pm 1.5 \times 10^{-3}$ |
| $R_g$ (Å) | 53<br>$\pm 0.13$ | 52<br>$\pm 0.12$ | 51<br>$\pm 0.13$ | 50<br>$\pm 0.19$ | 45<br>$\pm 0.20$ | 93<br>$\pm 2.05$ | 183<br>$\pm 3.9$ |
| $D_{max}$ (Å) | 190 | 188 | 180 | 185 | 154 | 350 | 650 |
| Porod volume (Å<sup>3</sup>) | 138000 | 141000 | 86000 | 78400 | 65000 | 268000 | 264000 |
| MW from Porod<br>volume (kDa) | 82 | 83 | 50 | 46 | 38 | 158 | 155 |
| $V_c$ (Å<sup>2</sup>) | 701 | 708 | 498 | 487 | 428 | 1030 | 1785 |
| MW from $V_c$ (kDa) | 82 | 83 | 43 | 40 | 36 | 101 | 155 |
| **DAMMIF *ab initio***<br>**modelling**<br>**(30 models)** |  |  |  |  |  |  |  |
| Symmetry | P22 | P22 | P1 | P1 | P1 | N/A | N/A |
| NSD mean and s.d. | 0.773<br>$\pm 0.120$ | 0.766<br>$\pm 0.184$ | 0.883<br>$\pm 0.040$ | 0.833<br>$\pm 0.031$ | 0.653<br>$\pm 0.016$ | N/A | N/A |
| $\chi^2$ (reference model) | 1.33 | 1.18 | 1.15 | 1.31 | 0.99 | N/A | N/A |
| **MONSA multi-phase *ab initio***<br>**modelling** |  |  |  |  |  |  |  |
| Symmetry | P2 | N/A | N/A | N/A | P2 | N/A | N/A |
| $\chi^2$ | 1.69 | N/A | N/A | N/A | 0.95-0.96 | N/A | N/A |
| **Structural modelling** |  |  |  |  |  |  |  |
| CRY SOL: Crystal<br>structure ($\chi^2$) | 178 | 200 | 32.2 | 14.2 | 2.32 | N/A | N/A |
| CRY SOL: + modelled C-terminal coiled-coil ($\chi^2$) | 13.5 | 20.2 | 8.84 | 3.40 | 1.23 | N/A | N/A |
| CORAL: + modelled<br>flexible N-termini ($\chi^2$) | 2.47 | 3.29 | 4.64 | 1.99 | N/A | N/A | N/A |
| FoXS: fibril model ($\chi^2$) | N/A | N/A | N/A | N/A | N/A | 1.22 | 1.09 |

### Crystal structure of SYCE2-TEX12 in a 4:4 assembly

We obtained an alternative crystal form of the SYCE2-TEX12 ΔCtip core complex, which diffracted anisotropically to 3.33-6.00 Å resolution, and was solved by molecular replacement using a three-helical bundle of the 2:2 structure as a search model (Table 1 and Supplementary Figure 4a). This revealed a 4:4 structure in which two 2:2 complexes interact laterally, through tessellation of S-shaped SYCE2 dimers, to create a molecule of the same length (14 nm) but twice as wide (4 nm) as individual 2:2 complexes (Figure 4a,b). The 2:2 crystal structure is largely retained by the two interacting 2:2 complexes (r.m.s. deviation = 1.91; Supplementary Figure 4b), which seamlessly generate an SYCE2 tetrameric core with staggered TEX12 chains arranged on opposing surfaces (Figure 4a). In common with the 2:2 structure, S2C sequences were missing from electron density, meaning that pairs of truncated SYCE2 and TEX12 C-termini are juxtaposed at either end of the 4:4 molecule (Figure 4c). The observed tessellation of 2:2 complexes provides an enticing mechanism for assembly of a 4:4 complex from structurally intact obligate 2:2 molecules, in agreement with our biochemical findings.

**Figure 4.**
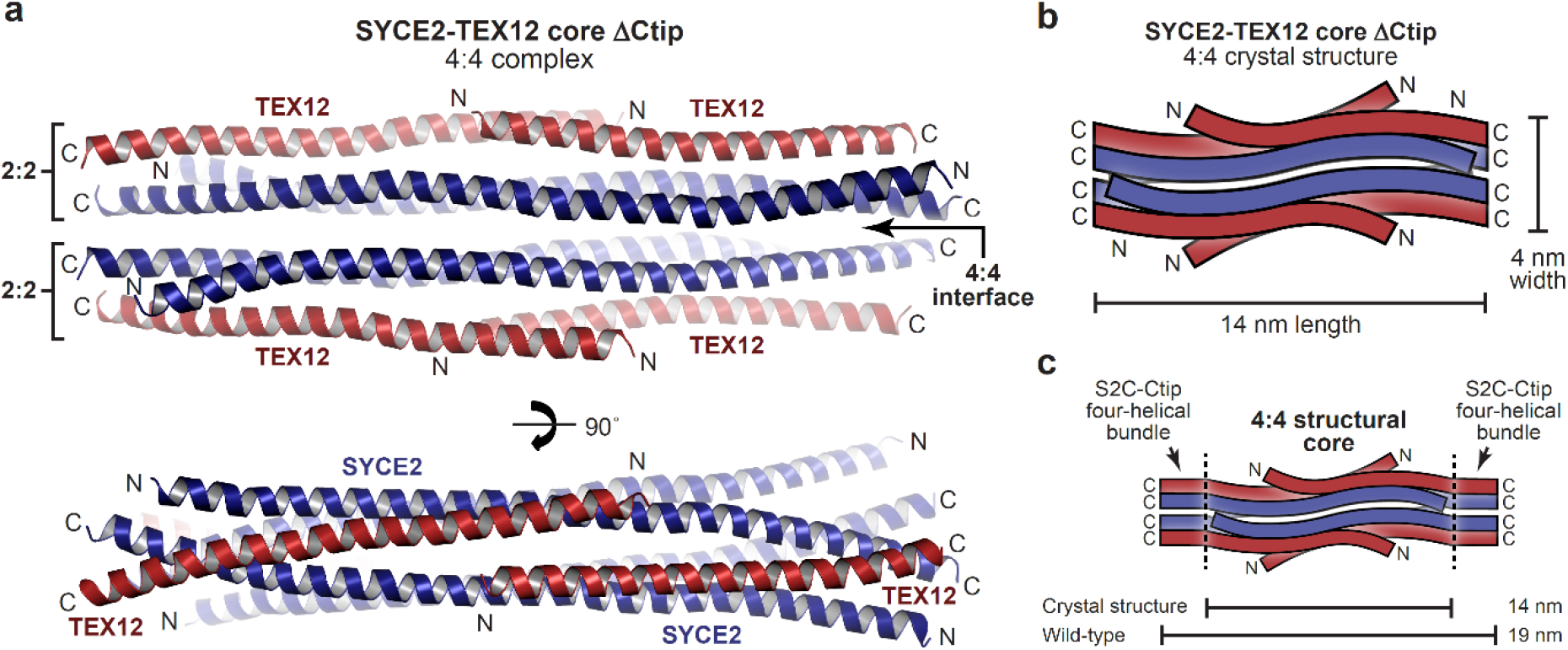
Crystal structure of the SYCE2-TEX12 core 4:4 complex. (**a**) Crystal structure of SYCE2-TEX12 core ΔCtip as a 4:4 complex. The structure consists of two 2:2 complexes interacting side-by-side through tessellation of their undulating SYCE2 surfaces at a 4:4 interface. This results in a central core of four SYCE2 chains (blue) flanked by two TEX12 chains (red) on either side. (**b-c**) Schematics of the 4:4 crystal structure in which (**b**) the orientation of SYCE2 and TEX12 chains are highlighted, and (**c**) their additional S2C and Ctip sequences are shown as C-terminal four-helical bundles. The 4:4 structure has a width of 4 nm and a length of 14 nm, which is predicted to increase to 19 nm upon addition of S2C and Ctip sequences.

### The SYCE2-TEX12 4:4 complex is stabilised by TEX12 C-terminal tip interactions

The 4:4 crystal structure lacks TEX12’s C-terminal tip, which we know is required for 4:4 assembly in solution, suggesting that this conformation was supported by the crystal lattice. We reasoned that the same tessellating 4:4 interface likely drives 4:4 assembly of the wild-type protein, which must be stabilised in solution by additional interactions of TEX12’s Ctip. To test this, we introduced glutamate mutations of amino-acids H89 and Y115, which mediate the tessellating interaction of SYCE2 dimers within the 4:4 crystal structure whilst having no structural role within constituent 2:2 complexes (Figure 5a). Within the core complex, the H89E Y115E mutation partially blocked 4:4 assembly, restricting 60% of material to a 2:2 complex, despite the presence of Ctip sequences (Figure 5b). SEC-SAXS determined that the wild-type 4:4 complex has a 19-nm length and 1.9-nm cross-sectional radius (Figure 5c,d and Supplementary Figure 6b-d), matching the observed geometry of the 4:4 crystal structure in which the length of the 2:2 complex is retained and its width is doubled from 2 nm to 4 nm (Figure 4). Further, multi-phase SAXS *ab initio* modelling of the wild-type 4:4 structure placed ΔCtip 2:2 *ab initio* envelopes in parallel, consistent with their lateral tessellating interaction, with missing mass – corresponding to S2C and Ctip sequences – placed at either end of the molecule (χ^2^ = 1.69; Figure 5e). Thus, the lateral tessellating interaction of 2:2 complexes within the 4:4 crystal structure describes the underlying architecture of the wild-type 4:4 assembly.

**Figure 5.**
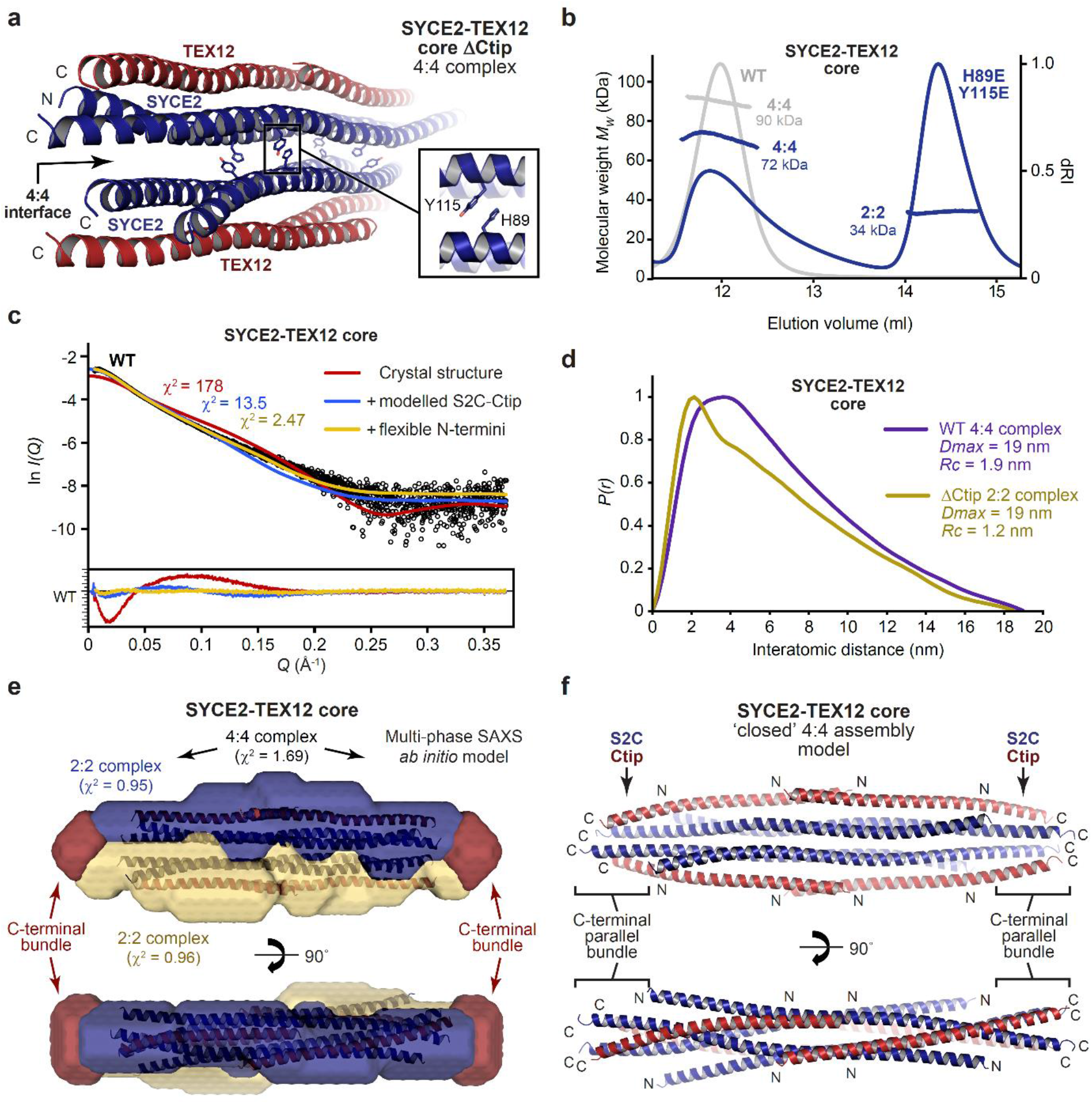
A molecular model of the SYCE2-TEX12 core ‘closed’ 4:4 assembly. (**a**) Crystal structure of the SYCE2-TEX12 core ΔCtip 4:4 complex, highlighting interactions between SYCE2 residues H89 and Y115 at the 4:4 interface between juxtaposed 2:2 complexes. (**b**) SEC-MALS analysis of SYCE2-TEX12 core H89E/Y115E showing the formation of a 34 kDa 2:2 complex (60%; theoretical – 44 kDa) and 72 kDa 4:4 complex (40%; theoretical – 89 kDa). SYCE2-TEX12 core wild-type is shown in grey for comparison. (**c-e**) SEC-SAXS analysis of SYCE2-TEX12 core. (**c**) SAXS scattering data of SYCE2-TEX12 core overlaid with the theoretical scattering curves of the 4:4 crystal structure (red), with modelled Ctip-S2C C-terminal bundles (blue), and with flexibly modelled N-termini (yellow); χ^2^ values are indicated and residuals for each fit are shown (inset). (**d**) SAXS *P(r)* interatomic distance distributions of SYCE2-TEX12 core wild-type and ΔCtip, showing maximum dimensions *(Dmax)* of 19 nm. Their cross-sectional radii (*Rc*) were determined as 1.9 nm and 1.2 nm, respectively. (**e**) Multiphase SAXS *ab initio* (MONSA) model of the SYCE2-TEX12 core 4:4 complex (entire envelope; χ^2^=1.69), consisting of two SYCE2-TEX12 core ΔS2C/ΔCtip complexes (blue and yellow envelopes; χ^2^=0.95 and 0.96) and remaining mass corresponding to S2C-Ctip C-terminal bundles (red). (**f**) Theoretical model of the SYCE2-TEX12 core ‘closed’ 4:4 complex in which S2C-Ctip hetero-dimeric coiled-coil models were docked onto the 4:4 crystal structure and assembled into C-terminal four-helical bundles through iterative energy minimisation and geometry idealisation.

How does TEX12’s Ctip stabilise the wild-type 4:4 complex? The location of pairs of truncated TEX12 and SYCE2 C-termini at either end of the 4:4 complex suggests that Ctip and S2C sequences could interact in four-helical structures that clamp together tessellated 2:2 molecules. In support of this, deletion of SYCE2’s S2C blocked 4:4 assembly and restricted the SYCE2-TEX12 core to a 2:2 complex (Supplementary Figure 5a), confirming that S2C and Ctip have equivalent roles in assembly. We noticed that the predicted heptad patterns of S2C-Ctip sequences are in-phase with the observed heptads of upstream SYCE2-TEX12 sequences at 2:2 ends of the crystal structure, so modelled an extended S2C-Ctip molecule as a parallel heterodimeric coiled-coil (Supplementary Figure 5b). This model docked seamlessly and in-phase with overlapping sequences within the 4:4 crystal structure (Supplementary Figure 5c), producing a model in which S2C-Ctip coiled-coils emanate from both ends of constituent 2:2 complexes (Supplementary Figure 5d). Cycles of energy minimisation and geometry idealisation allowed juxtaposed pairs of S2C-Ctip dimers to interact (Supplementary Figure 5d). This resulted in the formation of C-terminal bundles, consisting of parallel interactions between two S2C chains and two Ctip chains, which clamp together 2:2 complexes at either end of the tessellating 4:4 interface (Figure 5f). The resultant ‘clamped’ 4:4 structure closely fitted the SAXS scattering curve of the wild-type 4:4 core complex upon flexible modelling of missing N-termini (χ^2^ = 2.47; Figure 5c). Thus, formation of S2C-Ctip C-terminal bundles provides a mechanism for clamping together laterally associated 2:2 complexes within the architecture of the wild-type 4:4 complex.

### Controlling the hierarchical assembly of SYCE2-TEX12

Our model of the clamped 4:4 assembly allowed us to predict the sequence determinants of higher-order assembly by TEX12’s Ctip. The C-terminal bundle includes TEX12 amino-acids L110, F114, I117 and L121 in its hydrophobic core and F102, V116 and F109 in solvent-exposed positions (Figure 6a,b). We thus predicted that the first group is essential for clamped 4:4 assembly, whilst the second group is likely dispensable. Accordingly, hydrophobic core mutation L110E F114E I117E L121E (LFIL) restricted core and full-length SYCE2-TEX12 to 2:2 complexes and blocked fibrous assembly, thereby mimicking the Ctip deletion (Figure 6c-f). We similarly observed a restricted 2:2 complex upon analogous mutation of S2C hydrophobic core amino-acids V149, V153, V156 and L160 (Figure 6a and Supplementary Figure 6a). In contrast, surface mutation F102A V116A F109A (FVF) retained the 4:4 core complex, and wild-type pattern of full-length assembly in solution, but blocked higher-order assembly into fibres (Figures 2c,f and 6c-f). Importantly, SEC-SAXS confirmed that the 2:2 and 4:4 core complexes formed by LFIL and FVF mutants adopt the structures described for the ΔCtip and wild-type core complexes, respectively (Supplementary Figure 6b-g). We have thus identified separation-of-function mutations that block SYCE2-TEX12 in its building-block 2:2 structure and intermediate 4:4 assembly. The mimicry of Ctip deletion by LFIL mutation provides experimental validation for our proposed clamped 4:4 assembly model, whilst blockade of higher-order assembly by FVF mutation implicates these surface-exposed residues in the SYCE2-TEX12 fibre assembly mechanism.

**Figure 6.**
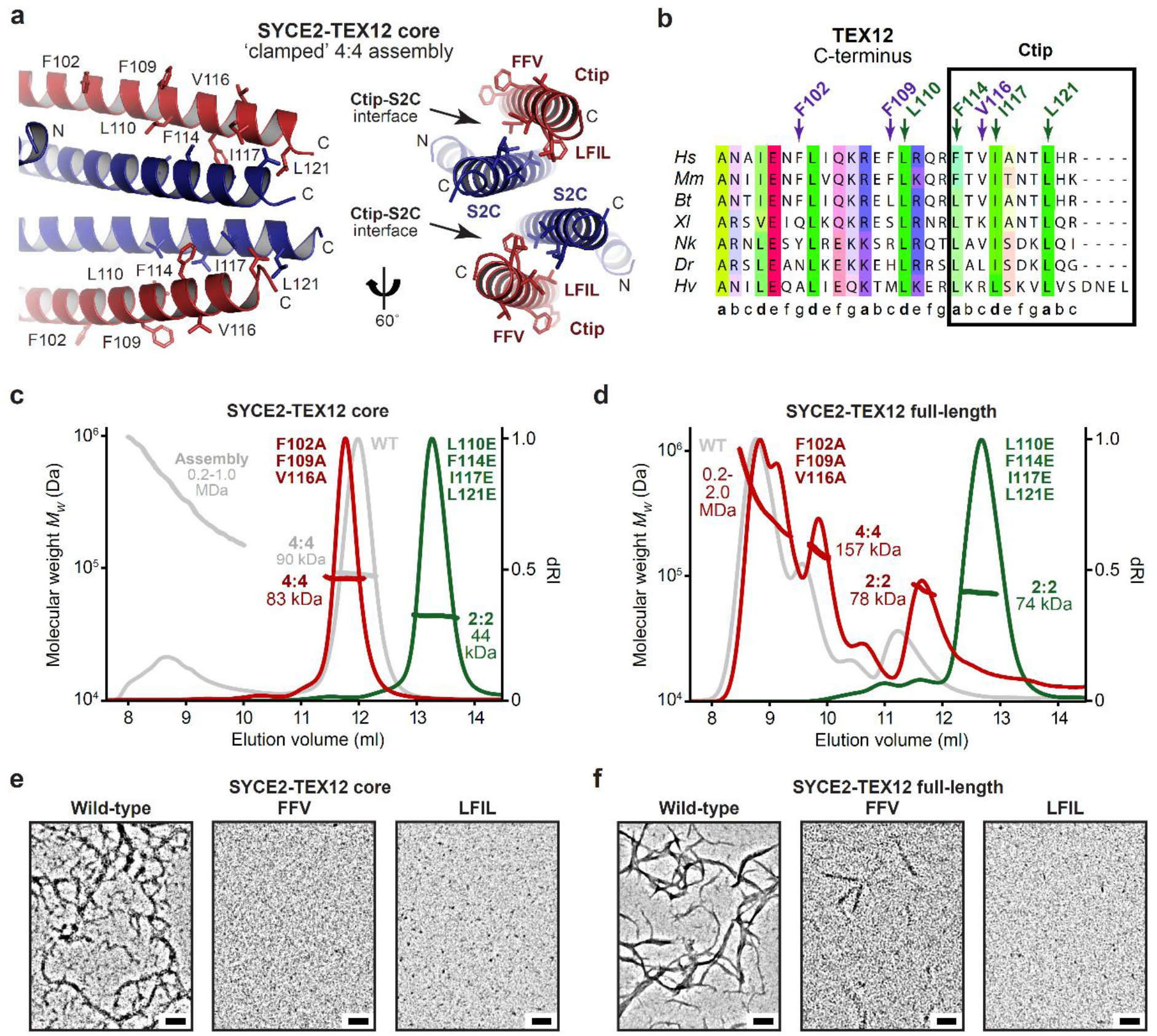
Molecular determinants of SYCE2-TEX12 self-assembly by TEX12’s C-terminal tip. (**a**) Molecular model of the SYCE2-TEX12 core ΔCtip ‘closed’ 4:4 assembly. The C-terminal bundle is formed of interactions between two S2C-Ctip coiled-coils, each of which involves heptad interactions between S2C residues V149, V153, V156 and L160, and Ctip residues L110, F114, I117 and L121, whilst F102, F109 and V116 are solvent-exposed. (**b**) Multiple sequence alignment of the TEX12 C-terminus, highlighting the Ctip sequence and the presence of LFIL residues at heptad positions, and FFV residues at non-heptad positions. (**c-d**) SEC-MALS analysis of SYCE2-TEX12 (**c**) core and (**d**) full-length mutants F102A F109A V116A (FFV, red) and L110E F114E I117E L121E (LFIL, green), with wild-type shown in grey for comparison. (**c**) SYCE2-TEX12 core FFV forms an 83 kDa 4:4 complex (theoretical – 88 kDa) with no higher-order assemblies, whilst LFIL forms a 44 kDa 2:2 complex (theoretical – 45 kDa). (**d**) SYCE2-TEX12 full-length FFV forms a 78 kDa 2:2 complex (15%; theoretical – 78 kDa), 155 kDa 4:4 complex (20%; theoretical – 156 kDa) and large molecular assemblies of up to 2.0 MDa (65%), whilst LFIL forms a 74 kDa 2:2 complex (theoretical – 78 kDa). (**e-f**) Electron microscopy of SYCE2-TEX12 (**e**) core and (**f**) full-length mutants. FFV inhibits SYCE2-TEX12 core assembly and restricts full-length to occasional fibres. LFIL inhibits SYCE2-TEX12 core and full-length fibrous assembly. Scale bars, 100 nm.

### SYCE2-TEX12 forms 2-nm and 4-nm fibres

How does TEX12’s Ctip drive assembly of 4:4 complexes into SYCE2-TEX12 fibres? The crystal lattice of the 4:4 structure is a fibrous array in which 4:4 molecules are arranged in end-to-end chains, with 1-nm gaps between their juxtaposed C-termini, and with a lateral stagger (Figure 7a). Could this represent the fibrous structure of SYCE2-TEX12, with end-to-end chains corresponding to fibres of 4:4 molecules that are thickened by lateral staggering? This would explain the stabilisation of constituent 4:4 complexes in absence of Ctip sequences. We obtained protein crystals of various SYCE2-TEX12 core constructs, including ΔCtip, which displayed a fibrous pattern of X-ray diffraction that is typical of the k-m-e-f family (keratin, myosin, epidermin, fibrinogen) of fibrous α-proteins (Figure 7b)^49–52^. This consists of a 5.1 Å meridional arc arising from the coiled-coil repeat, and 12 Å equatorial reflections that correspond to inter-helical distances of three-/four-helical bundles, in contrast with the classic 9.8 Å inter-helical distances of dimeric coiled-coils^49–52^. This is consistent with the end-to-end chains of molecules observed within the 4:4 crystal lattice. Further, upon rotation about the meridional axis, equatorial reflections are resolved into orthogonal 15-nm and 5-nm regularly-spaced reflections, closely matching the repeating units of the 4:4 crystal lattice (Figure 7a,b). Thus, the fibrous diffraction pattern is consistent with the fibrous array observed within the 4:4 crystal lattice, demonstrating that this arrangement of molecules is commonly exhibited by SYCE2-TEX12 constructs.

**Figure 7.**
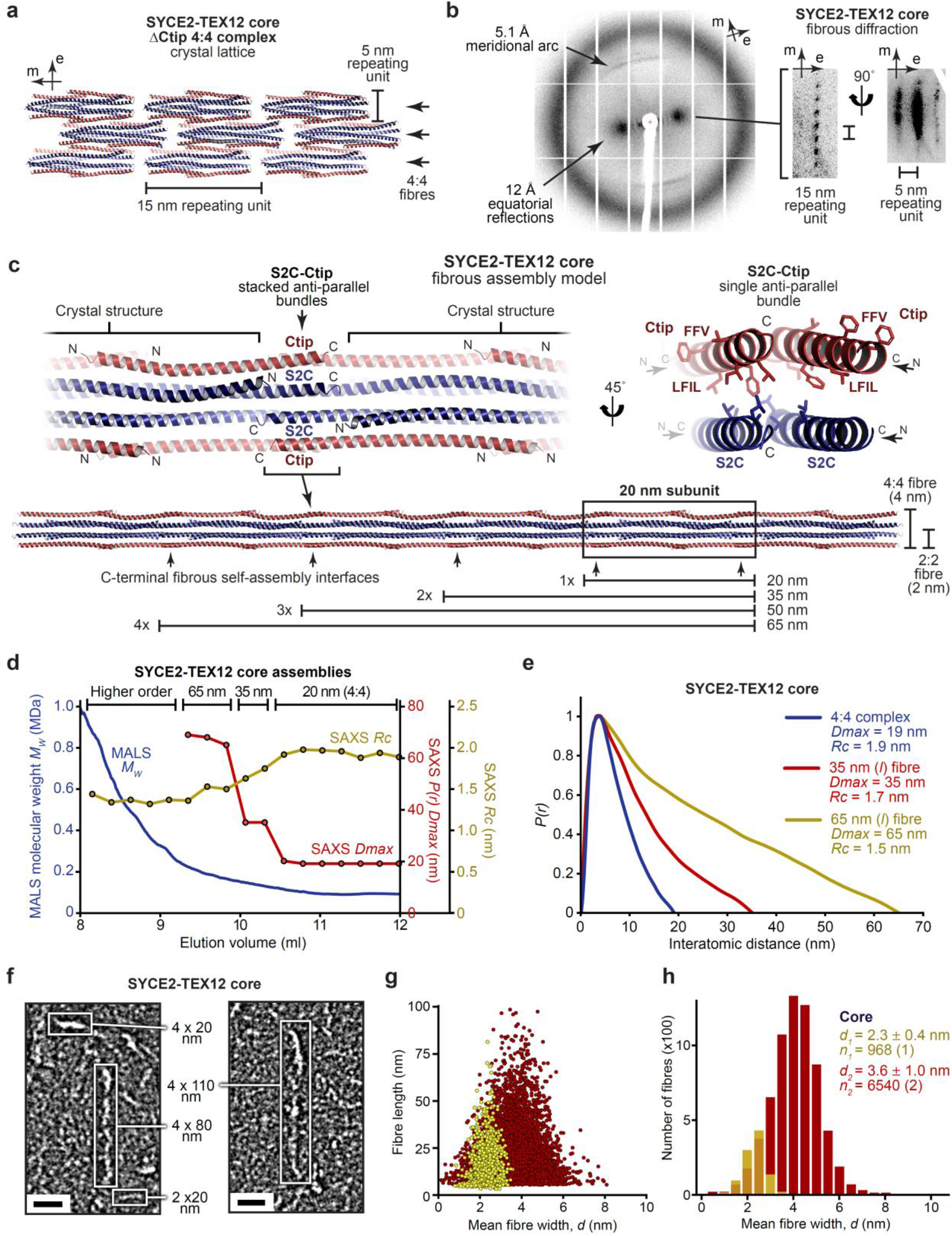
A molecular model of the SYCE2-TEX12 core fibrous assembly. (**a**) The SYCE2-TEX12 core ΔCtip 4:4 crystal lattice is formed of 15 nm translations along the meridional axis that generate 4:4 fibres, which associate laterally as 5 nm repeating units. (**b**) X-ray diffraction pattern of SYCE2-TEX12 core ΔCtip crystals demonstrating 5.1 Å meridional arcs and 12 Å equatorial reflections, which resolve upon rotation about the meridional axis into longitudinal and lateral spacings corresponding to 15 nm and 5 nm repeating units, respectively. (**c**) Theoretical model of SYCE2-TEX12 core fibrous assembly in which adjacent 4:4 complexes are translated by 15 nm and interact back-to-back through stacked S2C-Ctip anti-parallel four-helical bundles (left), of which each bundle contains a hydrophobic core of CtIP LFIL and S2C residues, with FFV residues remaining solvent-exposed (right). This assembly mechanism generates fibres of discrete lengths corresponding to the 20 nm initial subunit and integer multiples of the 15 nm repeating unit, with a width of 2 nm or 4 nm if it is formed of 2:2 or 4:4 complexes, respectively. (**d-e**) SEC-SAXS analysis of SYCE2-TEX12 core assemblies. (**d**) SAXS *P(r)* maximum dimensions *(Dmax,* red), cross-sectional radii (*Rc*, yellow) and corresponding MALS molecular weights (blue) are shown across an elution peak; *Dmax* is not determined for higher-order species in which the Shannon limit is exceeded. (**e**) SAXS *P(r)* interatomic distance distributions of the SYCE2-TEX12 core 4:4 complex (blue) and its fibrous assemblies of lengths 35 nm (red) and 65 nm (yellow). (**f-h**) Electron microscopy of SYCE2-TEX12 core in which (**f**) electron micrographs reveal molecules of 2 x 20 nm, 4 x 20 nm, 4 x 80 nm and 4 x 110 nm (w x l). Scale bars, 20 nm. (**g**) Scatter plots of mean fibre width (*d*, nm) against fibre length (nm), and (**h**) histograms of the number of fibres with mean widths within 0.5-nm bins, for two distinct populations (2 nm, yellow; 4 nm, red). The mean, standard deviation and number of fibres in each population were determined from one and two micrographs, respectively.

If the 4:4 crystal lattice represents the true fibrous assembly, then it should be possible to rationalise S2C-Ctip’s structural role within this fibrous array. We thus modelled S2C and Ctip as ideal helical extensions of SYCE2 and TEX12 chains within 4:4 complexes *in situ.* These ideal helical extensions immediately tessellated and formed favourable interactions between chains of end-to-end associated molecules within the crystal lattice (Figure 7c and Supplementary Figure 7a). This resulted in the formation of anti-parallel S2C-Ctip bundles between juxtaposed 2:2 ends, such that end-to-end interactions of 4:4 molecules are mediated by two stacked S2C-Ctip bundles (Figure 7c). Importantly, LFIL and S2C heptad residues are located within the hydrophobic core, whilst FVF residues are surface-exposed, suggesting roles in end-to-end and lateral fibre interactions, respectively, in agreement with the clamped 4:4 model and their mutant phenotypes (Figure 7c). Our molecular model suggests that constituent SYCE2-TEX12 fibres could be 2-nm or 4-nm wide, formed from 2:2 or 4:4 building-blocks, and stabilised by individual S2C-Ctip bundles or the tessellating 4:4 interface with stacked bundles (Figure 7c). Further, it predicts that the lengths of constituent fibres are defined by a 20-nm initial subunit with 15-nm increments for additional subunits (Figure 7c).

We tested the fibre assembly model through SAXS analysis of the molecular length and cross-sectional radius of species across the size-exclusion chromatography elution profile of SYCE2-TEX12 core (Figure 7d). There was a stepwise increase in molecular length from 19 nm (4:4) to 35 nm and 65 nm at elution points preceding the 4:4 complex, with a concomitant reduction in cross-sectional radius from 1.9 nm to 1.4 nm, suggesting a progression from pure 4:4 fibres (1.9-nm radius; 4-nm width) to a mixture with 2:2 fibres (1.2-nm radius; 2-nm width) (Figure 7d,e and Table 2). The 35-nm and 65-nm species correspond to the predicted lengths of end-to-end assemblies, and their SAXS scattering curves were fitted by modelled fibres of two 4:4 complexes (4 x 35 nm; χ^2^ = 1.22) and four 2:2 complexes (2 x 65 nm; χ^2^ = 1.09), respectively (Table 2 and Supplementary Figure 7b-e). We confirmed these findings by visualising the smallest SYCE2-TEX12 core assemblies within negatively-stained regions of electron micrographs (Figure 7f). Automated image analysis determined populations with mean fibre widths of 2.3 ± 0.4 nm and 3.6 ± 1.0 nm, corresponding to 2:2 (2-nm width) and 4:4 (4-nm width) fibres, respectively (Figure 7f-h). Further, the observed lengths of 2-nm and 4-nm fibres conformed to the predicted discrete lengths of end-to-end assemblies (Figure 7f). Thus, SAXS and EM data support our molecular model of SYCE2-TEX12 assembly into 2-nm and 4-nm fibres.

### Hierarchical assembly of SYCE2-TEX12 fibres

How do 2-nm and 4-nm fibres assemble into the 40-nm fibres that are readily formed by SYCE2-TEX12 and define the SC central element? We visualised full-length SYCE2-TEX12 within negatively-stained regions of electron micrographs (Figure 8a). This revealed the presence of 2-nm (2.1 ± 0.8 nm) and 4-nm (4.1 ± 1.6 nm) fibres, corresponding to the 2:2 and 4:4 fibres observed for the core complex, but of considerably greater length (Figure 8a-c). Importantly, the full-length complex also formed 10-nm fibres (9.6 ± 2.4 nm), which frequently intertwined within bundles of up to 40 nm that accumulated stain (Figure 8a-c). The 10-nm fibres likely represent laterally-associated 4:4 fibres, interacting through Ctip FVF residues and flanking termini, and seemingly represent an upper limit for the width of individual ‘smooth’ fibres (Figure 8d,e). The 40-nm bundled fibres correspond to the dimensions and polymorphic appearance of positively-stained SYCE2-TEX12 fibres (Figure 2a,d), and of the SC central element (Figure 1a), so likely represent an underlying architecture that is normally shrouded by accumulated stain. Thus, we conclude that SYCE2-TEX12 undergoes fibrous assembly through a hierarchical mechanism in which building-block 2:2 molecules laterally tesselate into 4:4 molecules, and assemble via end-to-end S2C-Ctip bundles to form 2-nm and 4-nm fibres (Figure 8d,e). These structures thicken through lateral interactions to form 10-nm fibres, which become interwoven into 40-nm bundled fibres (Figure 8d,e). Thus, we define the molecular mechanism whereby SYCE2-TEX12 self-assembles into rope-like fibres that structurally underpin longitudinal growth of the SC to enable synaptic elongation along the axial length of meiotic chromosomes.

**Figure 8.**
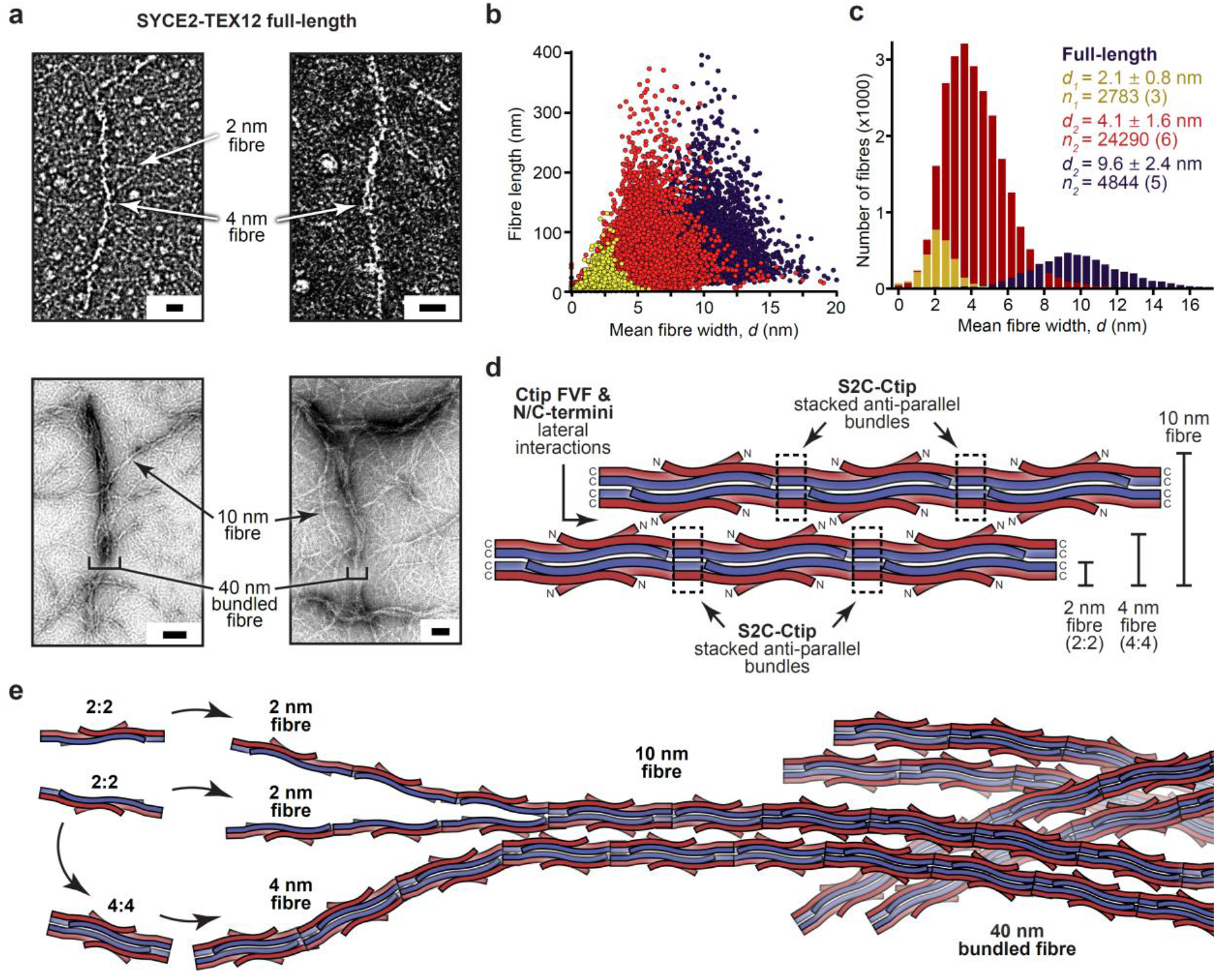
A molecular mechanism for fibrous assembly of SYCE2-TEX12. (**a-c**) Electron microscopy of full-length SYCE2-TEX12 in which (**a**) electron micrographs reveal the presence of 2 nm, 4 nm and 10 nm fibres, which become intertwined in bundled fibres of up to 40 nm in width. Scale bars, 20 nm (top) and 50 nm (bottom). (**b**) Scatter plots of mean fibre width (*d*, nm) against fibre length (nm), and (**c**) histograms of the number of fibres with mean widths within 0.5 nm bins, for three distinct populations (2 nm, yellow; 4 nm, red; 10 nm, blue). The mean, standard deviation and number of fibres in each population were determined from three, six and five micrographs, respectively. (**d**) Schematic of SYCE2-TEX12 fibrous assembly in which stacked S2C-Ctip anti-parallel bundles mediate end-to-end associations between subunits that generate 2-nm fibres and 4-nm fibres from 2:2 and 4:4 complexes, respectively. 10-nm fibres form through the lateral association of 4:4 fibres, mediated by Ctip FVF and N/C-terminal interactions. (**e**) Model of hierarchical fibrous assembly by SYCE2-TEX12. The building-block 2:2 complexes can assemble directly into 2-nm fibres, which then associate into 4-nm fibres, or can aggregate into ‘closed’ 4:4 structures that directly assemble into 4-nm fibres. The lateral association of 4-nm fibres generates 10-nm fibres, which become intertwined in bundled fibres of up to 40 nm width and many micrometres in length, and fulfil the known dimensions of the SC central element.

## Discussion

The formation of supramolecular cellular structures through protein self-assembly is of fundamental importance to numerous physiological systems, including cytoskeletal and chromosomal structure, and pathological processes such as amyloid formation^53–55^. Self-assembling systems can be defined in terms of minimum building-blocks that generate a range of assembled structures through recursive interactions. In meiotic chromosome synapsis, the supramolecular SC is formed by three major self-assembling systems, consisting of a zipper-like SYCP1 lattice, chromosome-associated SYCP3 axes, and a midline SYCE2-TEX12 fibrous backbone^23,37,38^. Here, we report that SYCE2-TEX12 is formed of 2:2 hetero-tetrameric building-blocks, which assemble into fibres through short motifs at the C-termini of their constituent α-helices. The deletion of TEX12’s C-terminal tip blocked fibrous assembly and enabled X-ray crystallographic structure solution of its 2:2 building-block complex, revealing an unusual combination of three- and four-helical coiled-coils within a rod-like structure. We previously demonstrated that SYCP1 and SYCP3 consist of homo-tetrameric rod-like building-blocks that self-assemble via motifs at the N- and C-terminal ends of their α-helical cores^23,38^. Thus, an apparent theme in mammalian SC assembly is the presence of tetrameric building-blocks, of rod-like four-helical coiled-coils, which self-assemble through short motifs at the coiled-coil’s termini that may be disrupted to stabilise the non-assembled building-block state in solution.

The assembly motifs of SYCE2-TEX12 mediate self-assembly in two distinct modes. Firstly, parallel four-helical bundles between S2C and Ctip motifs clamp together the ends of laterally-tessellated 2:2 molecules in discrete 4:4 assembly intermediates. Secondly, anti-parallel associations of S2C and Ctip motifs, in individual and stacked four-helical bundles, mediate end-to-end interaction of 2:2 and 4:4 molecules within 2-nm and 4-nm fibres, respectively. These clamped and fibrous conformations are stabilised by heptad interactions of the same hydrophobic amino-acids of Ctip (LFIL residues) and S2C (Figure 8d,e). In contrast, Ctip’s surface-exposed FVF residues mediate staggered lateral interactions that enable thickening into 10-nm ‘smooth’ fibres that become intertwined within mature 40-nm bundled fibres (Figure 8d,e). There are clear parallels with SYCP1 and SYCP3, in which supramolecular assembly is mediated by anti-parallel four-helical bundle formation between N-terminal tips of bioriented SYCP1 molecules^23^, and end-to-end interactions of juxtaposed motifs within SYCP3 fibres^38,43^. However, their resultant assemblies are structurally and functionally distinct owing to subtle differences in how these common assembly principles are implemented.

There are several intriguing differences between the assembly characteristics of SYCE2-TEX12 fulllength and core complexes. The full-length protein shows a higher propensity for assembly and exists in a wide range of solution species, which includes 2:2 and 4:4 complexes, whereas the core protein forms almost entirely 4:4 complexes. This suggests that unstructured termini of SYCE2 and TEX12 provide additional longitudinal stability to nascent assemblies that are formed through lateral and end-to-end interfaces. In their absence, a compensatory increase in the number of end-to-ends within the cross-section, through fibre thickening, may provide the additional cooperativity necessary to achieve structural stability, explaining our EM finding that core fibres are slightly thicker than fulllength fibres. Further, unstructured termini could stabilise 2:2 complexes by acting in ‘cis’ through termini interacting in the same manner with their own core, explaining the formation of wild-type 2:2 complexes by full-length but not core SYCE2-TEX12. Thus, the core complex may represent a restricted form of SYCE2-TEX12 in which the absence of unstructured termini reduces the propensity for higher-order assembly and also reduces the stability of building-block 2:2 complexes, resulting in its stabilisation within 4:4 complexes that represent partially assembled SYCE2-TEX12 structures.

The X-ray diffraction properties of SYCE2-TEX12 fibres are characteristic of the k-m-e-f family of fibrous α-proteins (keratin, myosin, epidermin and fibrinogen)^49–52^. These consist of 5.1 Å meridional arcs that arise from coiled-coil repeats oriented along the fibrous axis, and equatorial reflections that arise from lateral inter-helical spacings, which are 9.5-9.8 Å for dimeric coiled-coils, and 12 Å for SYCE2-TEX12’s three- and four-helical bundles^49–52^. Intermediate filaments (IF; including vimentin, lamin and keratin) constitute a subgroup of fibrous α-proteins that assemble from dimeric coiled-coil building-blocks, which associate along their long axis into tetramers that interact end-to-end and laterally within 10-nm fibres^56–62^. These stages of stepwise assembly appear analogous to SYCE2-TEX12’s 2:2 building-blocks, 4:4 assembly intermediates and 10-nm fibres. Further, IF proteins consist of coiled-coil cores flanked by assembly motifs^63^, their 10-nm fibres can assemble into meshwork and bundled networks^64,65^, and lamin forms 3.5-nm fibres that are reminiscent of SYCE2-TEX12’s 4-nm fibres^56,66^. Thus, hierarchical assembly of SYCE2-TEX12 bears striking resemblance to IF assembly, suggesting that SYCE2-TEX12 fibres may acquire the longitudinal strength, with flexion and torsional freedom, that is typical of the cytoskeleton by exploiting the same structural principles as molecules such as keratin.

What is the function of SYCE2-TEX12 in meiotic chromosome synapsis by the SC? The knockout phenotypes of SC central element proteins have suggested their grouping into initiation factors SYCE1, SYCE3 and SIX6OS1 that mediate synaptic initiation^28,30,32^, and elongation factors SYCE2 and TEX12 that extend synapsis along the chromosome length^33,34^. The hierarchical fibrous assembly of SYCE2-TEX12 provides structural rationale for this essential role in synaptic elongation. The formation of an IF-like SYCE2-TEX12 fibre at a site of nascent synapsis would provide a single longitudinal axis to guide and structurally underpin the extension of a single continuous synaptic event between each pair of homologous chromosomes. Thus, we propose that SYCE2-TEX12 provides a fibrous ‘backbone’ for the SC that defines a single continuous midline axis and underpins its long-range stability along the 4-24 μm length of meiotic chromosomes. This explains the presence of only short discontinuous stretches of nascent synapsis upon disruption of either SYCE2 or TEX1^33,34^. The supramolecular structures formed by SYCP1 and SYCP3 have distinct roles in physically tethering axes and stabilising compacted looped chromatin structure, respectively^23,38,39,43^. Thus, we conclude that the SC’s three major self-assembling systems – SYCP1, SYCP3 and SYCE2-TEX12 – adopt distinct and characteristic structures that fulfil specialised aspects of synaptic function, and the combination of these supramolecular structures achieves the enigmatic structure and function of the SC in meiosis.

## Materials and Methods

### Recombinant protein expression and purification

Sequences corresponding to regions of human SYCE2 (full-length, 1-218; core, 57-165; ΔS2C, 57-154) and TEX12 (full-length, 1-123; ΔCtip, 1-113; core, 49-123; core ΔCtip, 49-113) were cloned into pRSF-Duet1 (Novagen^®^) expression vectors for expression as TEV-cleavable N-terminal MBP- and His6-fusion proteins, respectively. Constructs were co-expressed in BL21 (DE3) cells (Novagen^®^), in 2xYT media, induced with 0.5 mM IPTG for 16 hours at 25°C. Cells were lysed by sonication in 20 mM Tris pH 8.0, 500 mM KCl, and fusion proteins were purified from clarified lysate through consecutive Ni-NTA (Qiagen), amylose (NEB) and HiTrap Q HP (GE Healthcare) ion exchange chromatography. Affinity tags were removed by incubation with TEV protease and cleaved samples were purified by HiTrap Q HP ion exchange chromatography and size exclusion chromatography (HiLoad™ 16/600 Superdex 200, GE Healthcare) in 20 mM Tris pH 8.0, 150 mM KCl, 2 mM DTT. Protein samples were concentrated using Pall Microsep™ Advance centrifugal devices, and were stored at −80°C following flash-freezing in liquid nitrogen. Protein samples were analysed by SDS-PAGE with Coomassie staining, and concentrations were determined by UV spectroscopy using a Cary 60 UV spectrophotometer (Agilent) with extinction coefficients and molecular weights calculated by ProtParam (http://web.expasy.org/protparam/).

### Crystallisation and structure solution of the SYCE2-TEX12 core ΔCtip 2:2 complex

SYCE2-TEX12 core ΔCtip protein crystals were obtained through vapour diffusion in sitting drops, by mixing 100 nl of 33 mg/ml protein at with 100 nl of crystallisation solution (35 % MPD) and equilibrating at 20°C for 4 months. Crystals were cryo-cooled in liquid nitrogen. X-ray diffraction data were collected at 0.9786 Å, 100 K, as 2000 consecutive 0.10° frames of 0.010 s exposure on a Pilatus3 6M detector at beamline I24 of the Diamond Light Source synchrotron facility (Oxfordshire, UK). Data were indexed, integrated in *XDS*^67^, scaled in *XSCALE*^68^, and merged with anisotropic correction and cut-off level at a local I/σ(I) of 1.2 using the *STARANISO* server^69^. Crystals belong to monoclinic spacegroup P21 (cell dimensions a = 88.52 Å, b = 24.19 Å, c = 88.48 Å, α = 90°, β = 115.7°, y = 90°), with a 2:2 SYCE2-TEX12 heterotetramer in the asymmetric unit. Structure solution was achieved through fragment-based molecular replacement using *ARCIMBOLDO_LITE*^70^, in which ten helices of 18 amino acids were placed by *PHASER*^71^ and extended by tracing in *SHELXE* utilising its coiled-coil mode^48^. A correct solution was identified by a *SHELXE* correlation coefficient of 47.8%. Model building was performed through iterative re-building by *PHENIX Autobuild*^72^ building in *COOT*^73^. The structure was refined using *PHENIX refine*^72^, using isotropic atomic displacement parameters with one TLS group. The structure was refined against data to anisotropy-corrected data with resolution limits between 2.42 Å and 3.36 Å, to *R* and *R_free_* values of 0.2871 and 0.3296 respectively, with 100% of residues within the favoured regions of the Ramachandran plot (0 outliers), clashscore of 6.89 and overall *MolProbity* score of 1.38^74^.

### Crystallisation and structure solution of the SYCE2-TEX12 core ΔCtip 4:4 complex

SYCE2-TEX12 core ΔCtip protein crystals were obtained through vapour diffusion in sitting drops, by mixing 100 nl of protein at 33 mg/ml with 100 nl of crystallisation solution (0.1 M sodium acetate, pH 4.0, 65 % MPD) and equilibrating at 20°C for 4 months. Crystals were cryo-cooled in liquid nitrogen. X-ray diffraction data were collected at 0.9688 Å, 100 K, as 2000 consecutive 0.10° frames of 0.020 s exposure on a Pilatus3 6M detector at beamline I24 of the Diamond Light Source synchrotron facility (Oxfordshire, UK). Data were processed using *AutoPROC*^75^, in which indexing, integration and scaling were performed by *XDS*^67^ and *XSCALE*^68^, and anisotropic correction with a local I/σ(I) cut-off of 1.2 was performed by *STARANISO*^69^. Crystals belong to orthorhombic spacegroup P21212 (cell dimensions a = 42.67 Å, b = 59.68 Å, c = 156.49 Å, α = 90°, β = 90°, γ = 90°), with a 2:2 SYCE2-TEX12 heterotetramer in the asymmetric unit. Structure solution was achieved by molecular replacement using *PHASER*^71^, in which two copies of one end of the higher resolution SYCE2-TEX12 2:2 structure (PDB accession 6R17) were placed in the asymmetric unit. The structure was completed through manual building in *COOT*^73^, with iterative refinement using *PHENIX refine*^72^. The structure was refined against data to anisotropy-corrected data with resolution limits between 3.33 Å and 6.00 Å, to *R* and *R_free_* values of 0.3266 and 0.3672 respectively, with 100% of residues within the favoured regions of the Ramachandran plot (0 outliers), clashscore of 3.48 and overall *MolProbity* score of 1.14^74^.

### Crystallisation and X-ray diffraction analysis of SYCE2-TEX12 core ΔCtip fibres

SYCE2-TEX12 core ΔCtip protein crystals were obtained through vapour diffusion in hanging drops, by mixing 1 μl of protein at 17 mg/ml in buffer containing 20% glycerol with 1 μl of crystallisation solution (0.05 M caesium chloride, 0.1 M MES pH 6.5, 30% v/v Jeffamine M-600) and equilibrating at 20°C. Crystals were cryo-cooled in liquid nitrogen in crystallisation solution supplemented with 25% glycerol. X-ray diffraction data were collected at 0.9763 Å, 100 K, as individual 0.50° frames of 0.500 s exposure on a Pilatus3 6M detector at beamline I03 of the Diamond Light Source synchrotron facility (Oxfordshire, UK). Diffraction images were analysed using *Adxv.*

### Size-exclusion chromatography multi-angle light scattering (SEC-MALS)

The absolute molecular masses of SYCE2-TEX12 complexes were determined by size-exclusion chromatography multi-angle light scattering (SEC-MALS). Protein samples at >1 mg/ml were loaded onto a Superdex™ 200 Increase 10/300 GL size exclusion chromatography column (GE Healthcare) in 20 mM Tris pH 8.0, 150 mM KCl, 2 mM DTT, at 0.5 ml/min using an ÄKTA™ Pure (GE Healthcare). The column outlet was fed into a DAWN^®^ HELEOS™ II MALS detector (Wyatt Technology), followed by an Optilab^®^ T-rEX™ differential refractometer (Wyatt Technology). Light scattering and differential refractive index data were collected and analysed using ASTRA^®^ 6 software (Wyatt Technology). Molecular weights and estimated errors were calculated across eluted peaks by extrapolation from Zimm plots using a dn/dc value of 0.1850 ml/g. SEC-MALS data are presented as differential refractive index (dRI) profiles, with fitted molecular weights (MW) plotted across elution peaks.

### Circular dichroism (CD) spectroscopy

Far UV circular dichroism (CD) spectroscopy data were collected on a Jasco J-810 spectropolarimeter (Institute for Cell and Molecular Biosciences, Newcastle University). CD spectra were recorded in 10mM Na_2_HPO_4_/ NaH_2_PO_4_ pH 7.5, at protein concentrations between 0.1-0.5 mg/ml, using a 0.2 mm pathlength quartz cuvette (Hellma), at 0.2 nm intervals between 260 and 185 nm at 4°C. Spectra were averaged across nine accumulations, corrected for buffer signal, smoothed and converted to mean residue ellipticity ([θ]) (x1000 deg.cm^2^.dmol^-1^.residue^-1^). Deconvolution was performed using the CDSSTR algorithm of the Dichroweb server (http://dichroweb.cryst.bbk.ac.uk)^76,77^. CD thermal denaturation was performed in 20 mM Tris pH 8.0, 150 mM KCl, 2 mM DTT, at protein concentrations between 0.1-0.4 mg/ml, using a 1 mm pathlength quartz cuvette (Hellma). Data were recorded at 222 nm, between 5°C and 95°C, at 0.5°C intervals with ramping rate of 2°C per minute, and were converted to mean residue ellipticity ([θ_222_]) and plotted as % unfolded ([θ]222,x-[θ]_222,5_)/([θ]_222,95-_[θ]_222,5_). Melting temperatures (Tm) were estimated as the points at which samples are 50% unfolded.

### Electron Microscopy

Electron microscopy (EM) was performed using FEI Philips CM100 and 120 kV Hitachi HT7800 transmission electron microscopes at the Electron Microscopy Research Services, Newcastle University. Protein samples at 0.005-3 mg/ml were applied to carbon-coated grids and negative staining was performed using 2% (weight/volume) uranyl acetate. Images were analysed in the Fiji distribution of *ImageJ*^78^, by applying background correction and an FFT Bandpass filter, and measuring the number, mean width and length of fibres using the Ridge detection algorithm.

### Size-exclusion chromatography small-angle X-ray scattering (SEC-SAXS)

SEC-SAXS experiments were performed at beamline B21 of the Diamond Light Source synchrotron facility (Oxfordshire, UK). Protein samples at concentrations >5 mg/ml were loaded onto a Superdex™ 200 Increase 10/300 GL size exclusion chromatography column (GE Healthcare) in 20 mM Tris pH 8.0, 150 mM KCl at 0.5 ml/min using an Agilent 1200 HPLC system. The column outlet was fed into the experimental cell, and SAXS data were recorded at 12.4 keV, detector distance 4.014 m, in 3.0 s frames. Data were subtracted and averaged, and analysed for Guinier region *Rg* and cross-sectional *Rg (Rc)* using ScÅtter 3.0 (http://www.bioisis.net), and *P(r)* distributions were fitted using *PRIMUS*^79^. *Ab initio* modelling was performed using *DAMMIF*^80^, in which 30 independent runs were performed in P1 or P22 symmetry and averaged. Multi-phase SAXS *ab initio* modelling was performed using *MONSA*^81^. Crystal structures and models were docked into *DAMFILT* and *MONSA* molecular envelopes using *SUPCOMB*^82^, and were fitted to experimental data using *CRYSOL*^83^ and *FoXS*^84^, and flexible termini were modelled and fitted to experimental data using *CORAL*^85^.

### Structural modelling

A heterodimeric coiled-coil of the SYCE2 and TEX12 C-termini, including their S2C and Ctip sequences (amino-acids 155-165 and 114-123, respectively) was modelled by *CCbuilder 2.0*^86^ and was docked onto the 2:2 and 4:4 crystal structures using *PyMOL* Molecular Graphics System, Version 2.3.2 Schrödinger, LLC. The 4:4 ‘closed’ assembly was modelled by manual editing of C-terminal helical positioning in *COOT*^73^, followed by iterations of energy minimisation using Rosetta Rela^x87^ interspersed with idealisation by PHENIX geometry minimisation^72^. The 4:4 ‘open’ fibre assembly was modelled by imposing the crystallographic translation symmetry of 156 Å along the long axis onto 4:4 complexes with modelled C-terminal coiled-coils, followed by manual editing of C-terminal helical positioning in *COOT*^73^, with iterations of energy minimisation using *Rosetta Relax*^87^ interspersed with idealisation by *PHENIX* geometry minimisation^72^.

### Protein sequence and structure analysis

Multiple sequence alignments were generated using *Jalview*^88^, and molecular structure images were generated using the *PyMOL* Molecular Graphics System, Version 2.0.4 Schrödinger, LLC.

### Accession codes and data availability

Crystallographic structure factors and atomic co-ordinates have been deposited in the Protein Data Bank (PDB) under accession numbers 6R17 and 6YQF. All other data are available from the corresponding author upon reasonable request.

## Acknowledgements

We thank Diamond Light Source and the staff of beamlines I03, I24 and B21 (proposals mx13587, mx18598, sm14435, sm15580, sm15897, sm15836 and sm21777). We thank I. Usón for advice on *ARCIMBOLDO_LITE,* and A. Baslé and H. Waller for assistance with X-ray crystallographic and CD data collection. O.R.D. is a Wellcome Trust Senior Research Fellow (Grant Number 219413/Z/19/Z).

## Author contributions

J.M.D. crystallised SYCE2-TEX12 2:2 and 4:4 complexes, and performed biophysical and electron microscopy experiments. L.J.S. collected initial biophysics data. O.R.D. solved the SYCE2-TEX12 crystal structures, analysed data, designed experiments and wrote the manuscript.

---

**Supplementary Figure 1.**
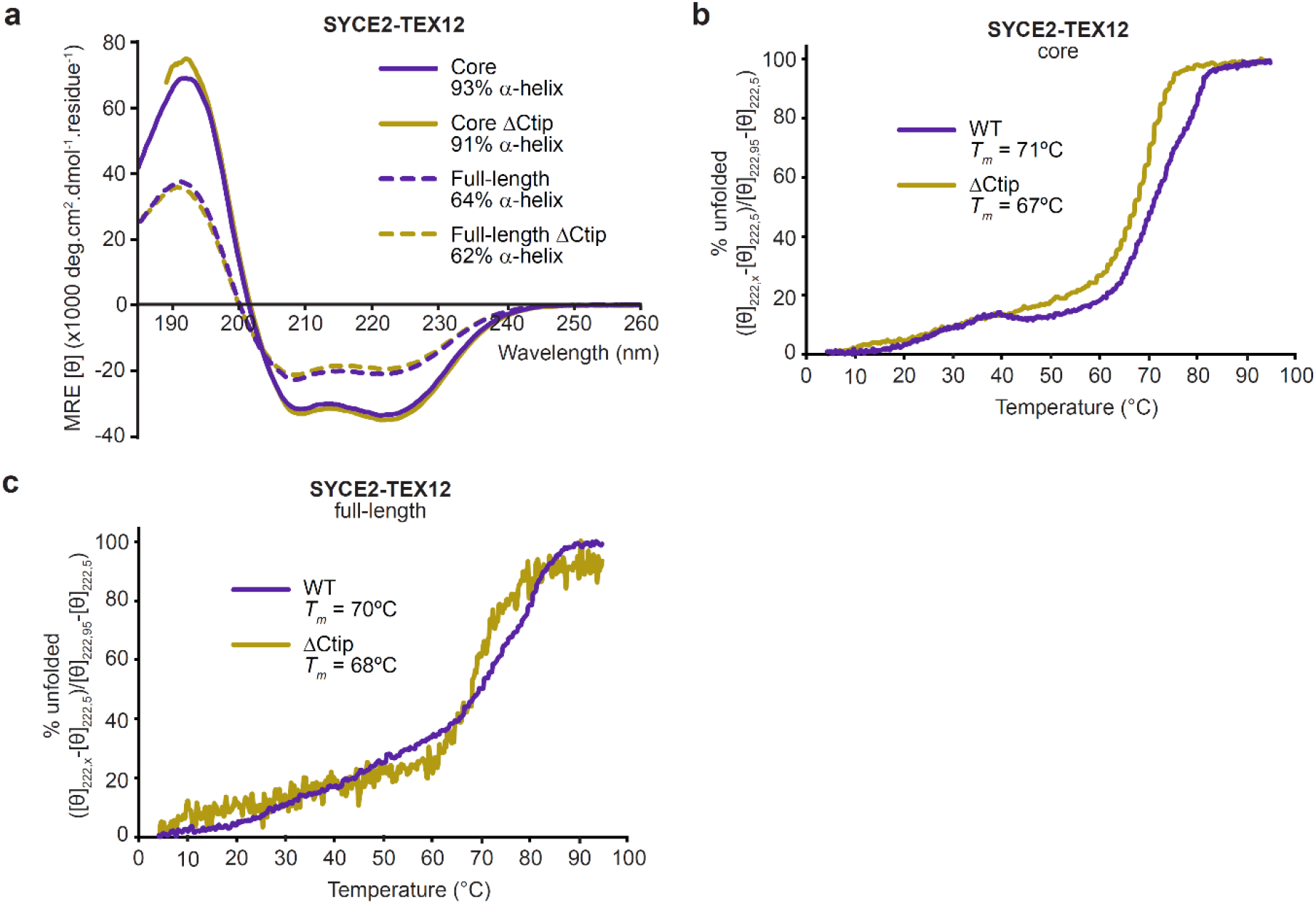
Circular dichroism analysis of SYCE2-TEX12 complexes. (**a-c**) Circular dichroism (CD) analysis of SYCE2-TEX12 core and full-length complexes containing wildtype (WT; purple) and ΔCtip (yellow) TEX12 sequences. (**a**) Far UV circular dichroism (CD) spectra recorded between 260 nm and 185 nm in mean residue ellipticity, MRE ([θ]) (x10^3^ deg.cm^2^.dmol^-1^.residue^-1^). Data were deconvoluted using the CDSSTR algorithm, with helical content indicated. (**b-c**) CD thermal denaturation of SYCE2-TEX12 (**b**) core and (**c**) full-length complexes, recording the CD helical signature at 222 nm between 5°C and 95°C, as % unfolded. Melting temperatures were estimated, as indicated.

**Supplementary Figure 2.**
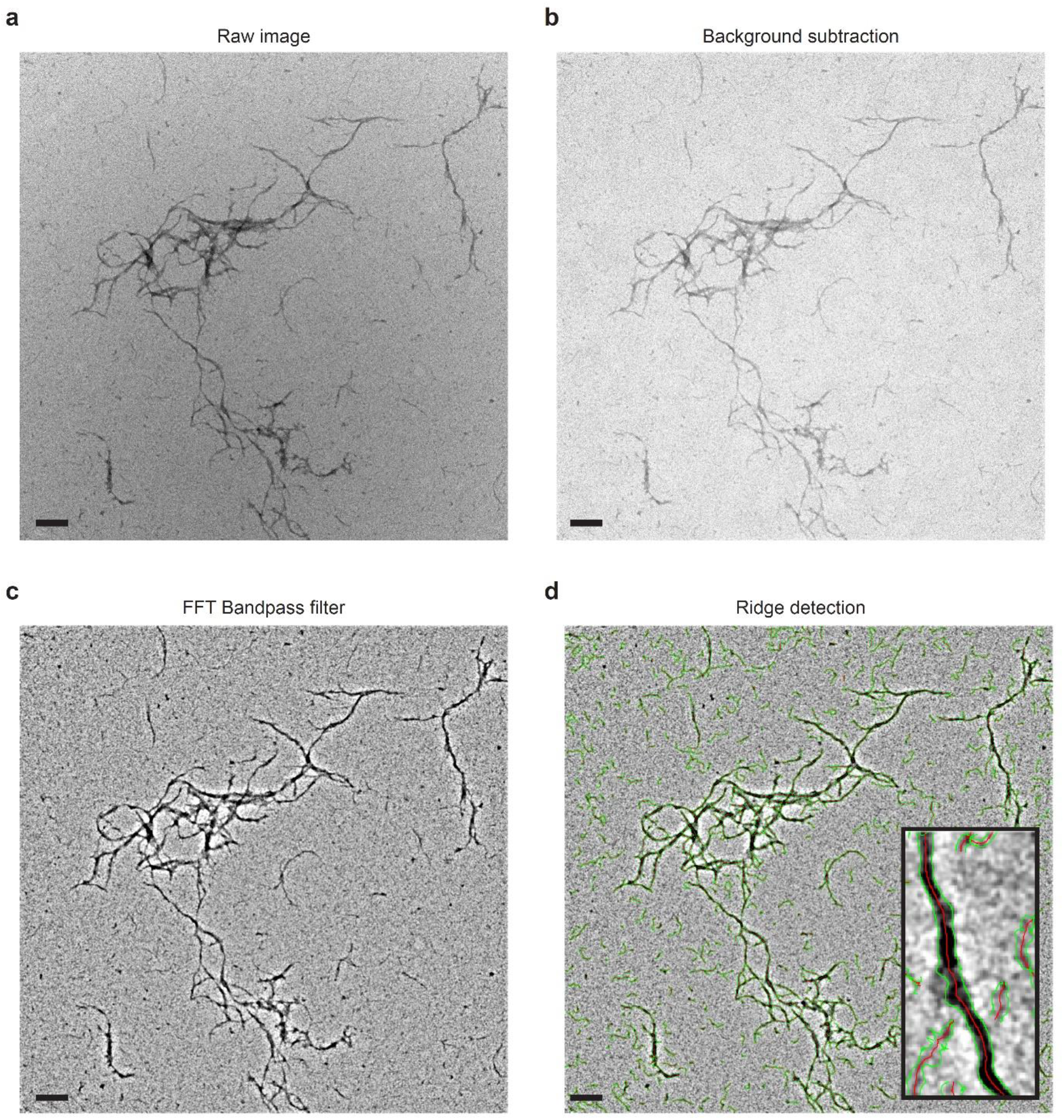
Image analysis of SYCE2-TEX12 electron micrographs. (**a-d**) The procedure used for automated image analysis and quantification of SYCE2-TEX12 fibres is shown for an example micrograph. The Fiji distribution of ImageJ was used to process (**a**) raw micrographs through (**b**) background subtraction and (**c**) FFT Bandpass filter. (**d**) Fibres were detected by the Ridge detection algorithm in which the centre and edges of interpreted fibres were highlighted in red and green, respectively, from which mean fibre widths and fibre lengths were determined.

**Supplementary Figure 3.**
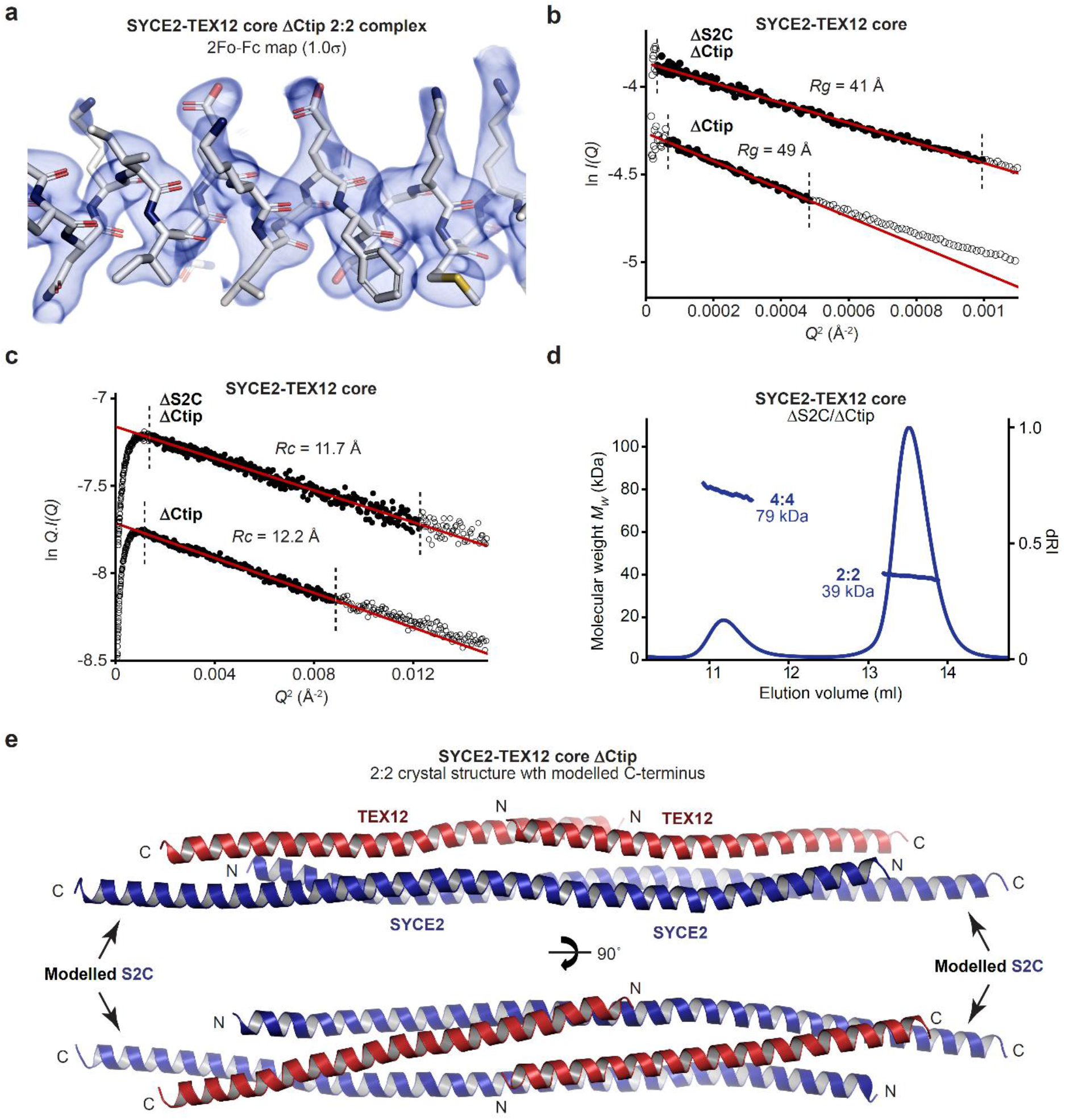
Crystal structure of the SYCE2-TEX12 core 2:2 complex. (**a**) 2Fo-Fc electron density map of the SYCE2-TEX12 core ΔCtip 2:2 complex (1.0σ) superimposed on the refined crystallographic model. (**b-c**) SEC-SAXS analysis of SYCE2-TEX12 core ΔS2C/ΔCtip. (**b**) SAXS Guinier analysis to determine the radius of gyration *(Rg);* linear fits are shown in red, with the fitted data range highlighted in black and demarcated by dashed lines. The *Q.Rg* values were < 1.3 and *Rg* was calculated as 41 Å and 49 Å, respectively. (**c**) SAXS Guinier analysis to determine the radius of gyration of the cross-section (*Rc*); linear fits are shown in red, with the fitted data range highlighted in black and demarcated by dashed lines. The *Q.Rc* values were < 1.3 and *Rc* was calculated as 11.7 Å and 12.2 Å, respectively. (**d**) SEC-MALS analysis of SYCE2-TEX12 core ΔS2C/ΔCtip showing the formation of a 39 kDa 2:2 complex (86%; theoretical – 40 kDa) and a 79 kDa 4:4 complex (14%; theoretical – 79 kDa). (**e**) Theoretical model of the full SYCE2-TEX12 core ΔCtip 2:2 complex in which the missing C-terminal coiled-coil and S2C helix were docked onto the crystal structure.

**Supplementary Figure 4.**
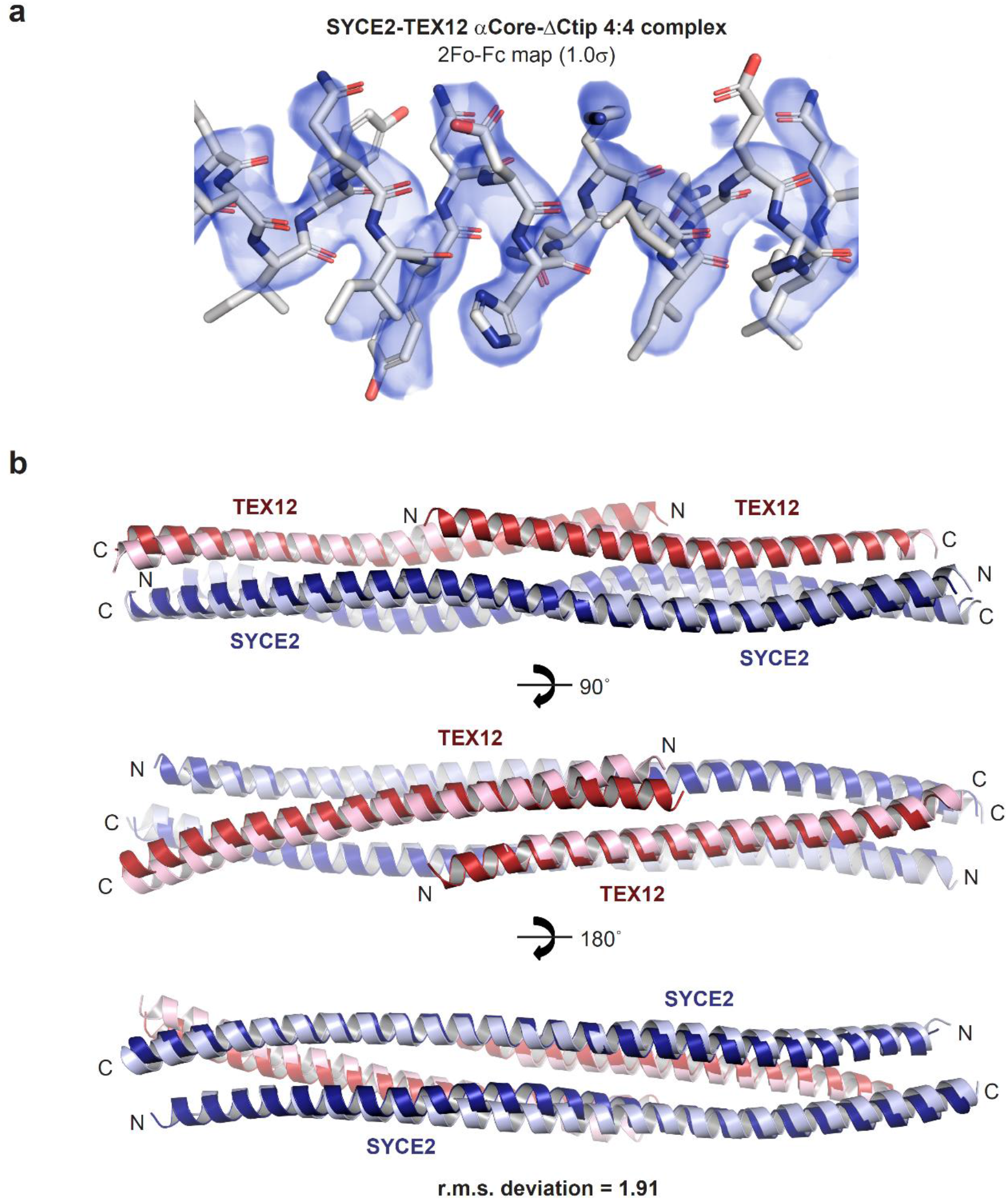
Crystal structure of the SYCE2-TEX12 core 4:4 complex. (**a**) 2Fo-Fc electron density map of the SYCE2-TEX12 core ΔCtip 4:4 complex (1.0σ) superimposed on the refined crystallographic model. (**b**) Superposition of a constituent 2:2 complex from the SYCE2- TEX12 core ΔCtip 4:4 structure (light blue and light red) and the SYCE2-TEX12 core ΔCtip 2:2 structure (dark blue and dark red), with r.m.s. deviation of 1.91.

**Supplementary Figure 5.**
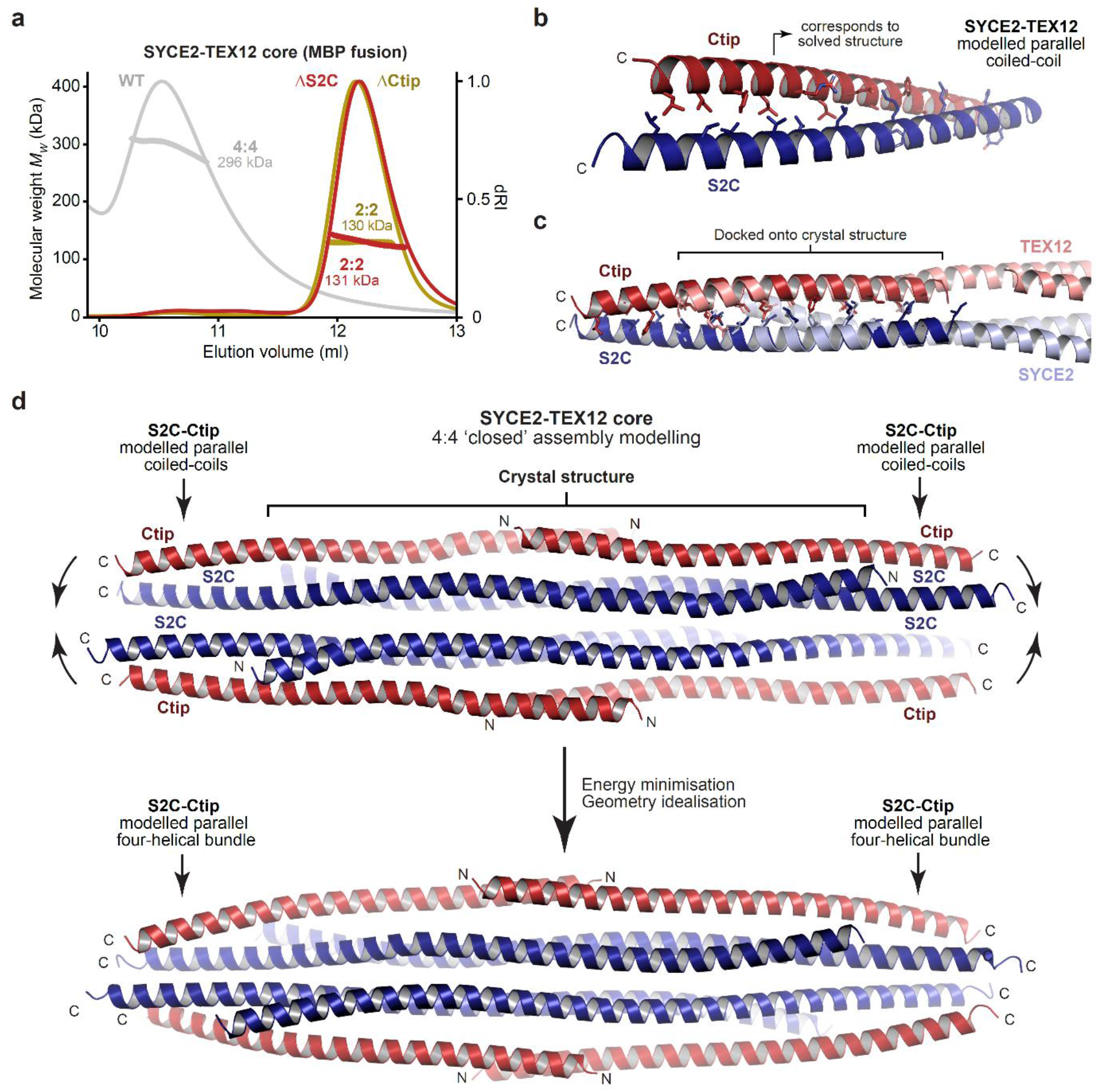
Modelling of the SYCE2-TEX12 core ‘closed’ 4:4 assembly. (**a**) SEC-MALS analysis of SYCE2-TEX12 core MBP-fusion proteins. SYCE2-TEX12 core wild-type is a 296 kDa 4:4 complex (theoretical – 278 kDa), whereas ΔS2C and ΔCtip form 131 kDa and 130 kDa 2:2 complexes, respectively (theoretical – 136 kDa and 137 kDa. (**b-d**) Modelling of the ‘closed’ 4:4 assembly. (**b**) Theoretical models of hetero-dimeric coiled-coils corresponding to SYCE2 and TEX12 amino-acids 114-165 and 75-123, respectively (including Ctip and S2C sequences 155-165 and 114123), were generated using *CCBuilder* by specifying the heptad register observed in the 2:2 and 4:4 crystal structures. (**c**) The coiled-coil models (red and blue) were docked onto the 2:2 ends of the 4:4 crystal structure (pale red and pale blue), showing close correspondence between overlapping helices. (**d**) This resulted in a 4:4 structure with emanating S2C-Ctip dimers (top) that was subjected to iterative energy minimisation and geometry idealisation such that S2C-Ctip sequences of adjacent dimers combined into capping four-helical bundles at either end of the 4:4 molecule (bottom).

**Supplementary Figure 6.**
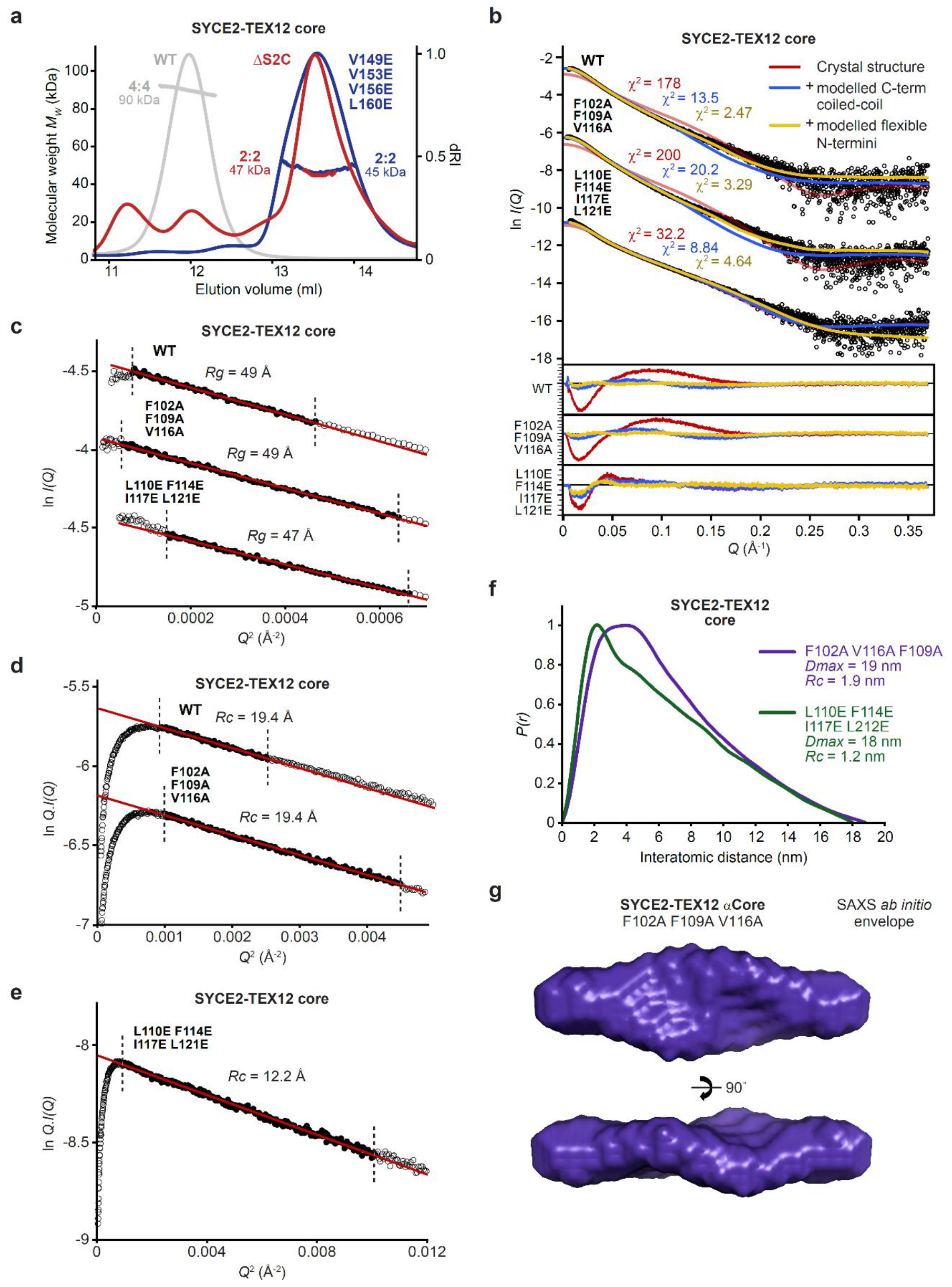
MALS and SAXS analysis of SYCE2-TEX12 wild-type and mutant complexes. (**a**) SEC-MALS analysis of SYCE2-TEX12 core ΔS2C and V149E, V153E, V156E and L160E mutants, demonstrating the formation of 47 kDa and 45 kDa 2:2 complexes, respectively (theoretical – 42 kDa and 45 kDa). The wild-type 90 kDa 4:4 complex is shown in grey for comparison. (**b-g**) SEC-SAXS analysis of SYCE2-TEX12 core wild-type, F102A F109A V116A (FFV) and L110E F114E I117E L121E (LFIL) mutants. (**b**) SAXS scattering data overlaid with the theoretical scattering curves of the 4:4 crystal structure (red), with modelled Ctip-S2C C-terminal bundles (blue), and with flexibly modelled N- termini (yellow); χ^2^ values are indicated and residuals for each fit are shown (inset). (**c**) SAXS Guinier analysis to determine the radius of gyration (*Rg*); linear fits are shown in red, with the fitted data range highlighted in black and demarcated by dashed lines. The *Q.Rg* values were < 1.3 and *Rg* was calculated as 49 Å, 49 Å and 47 Å, respectively. (**d-e**) SAXS Guinier analysis to determine the radius of gyration of the cross-section (*Rc*); linear fits are shown in red, with the fitted data range highlighted in black and demarcated by dashed lines. The *Q.Rc* values were < 1.3 and *Rc* was calculated as (**d**) 19.4 Å for wildtype and FFV and (**e**) 12.2 Å for LFIL. (**f**) SAXS *P(r)* interatomic distance distributions of SYCE2-TEX12 core FFV and LFIL, showing maximum dimensions of 19 nm and 18 nm, respectively. (**g**) SAXS *ab initio* model of SYCE2-TEX12 core FFV in which a filtered averaged model from 30 independent DAMMIF runs is shown.

**Supplementary Figure 7.**
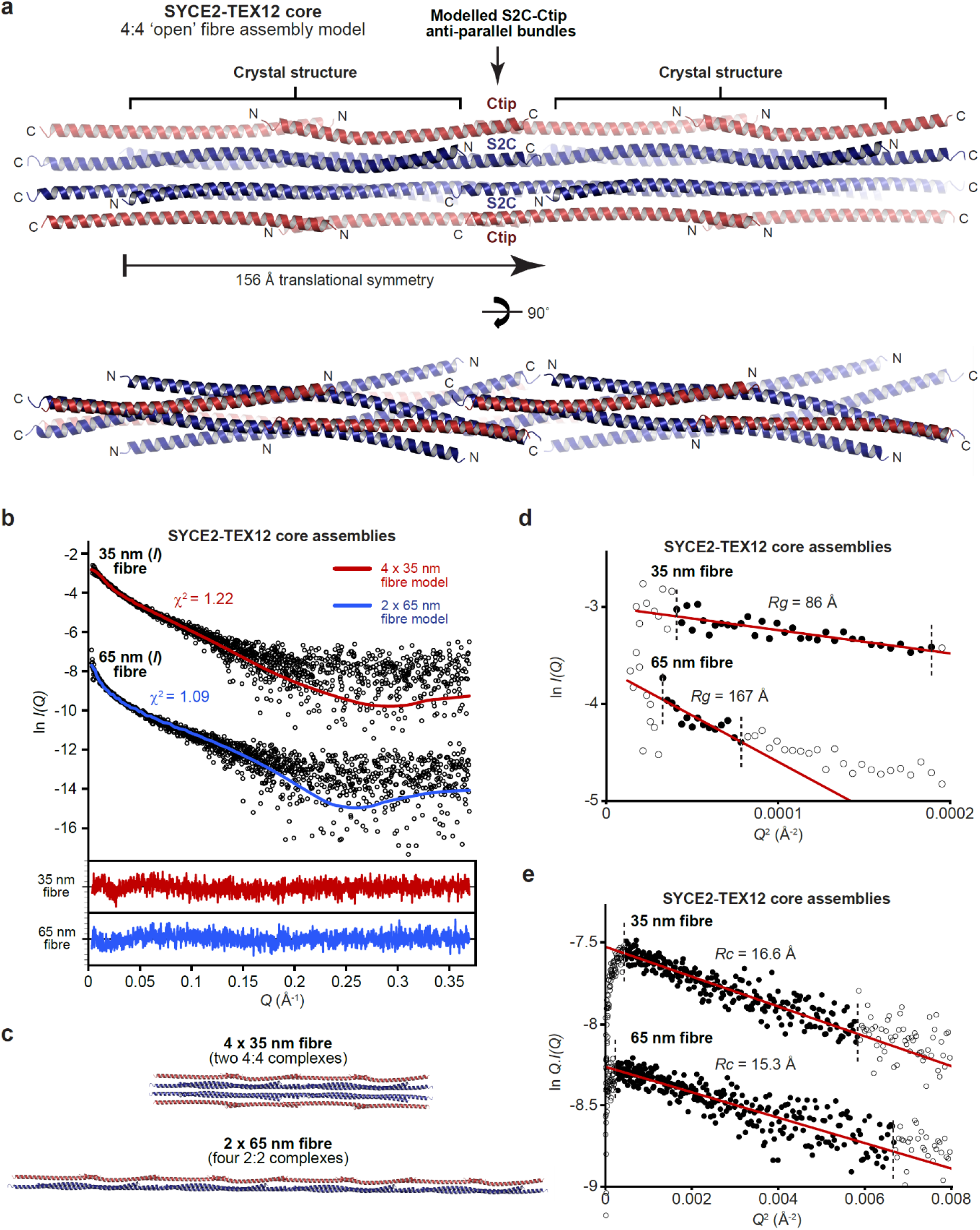
Modelling of the SYCE2-TEX12 core fibrous assembly. (**a**) Model of SYCE2-TEX12 core fibrous assembly in which adjacent 4:4 complexes are translated by 15 nm and interact back-to-back through stacked S2C-Ctip anti-parallel four-helical bundles. (**b-e**) SEC- SAXS analysis of SYCE2-TEX12 core 35 nm and 65 nm (length) fibres. (**b**) SAXS scattering data overlaid with the theoretical scattering curves of a 4 x 35 nm (w x l) fibre model (two consecutive 4:4 complexes, red) and a 2 x 65 nm (w x l) fibre model (four consecutive 2:2 complexes, blue); χ^2^ values are indicated and residuals for each fit are shown (inset). (**c**) The 4 x 35 nm and 2 x 65 nm fibre models used for SAXS data fitting. (**d**) SAXS Guinier analysis to determine the radius of gyration *(Rg);* linear fits are shown in red, with the fitted data range highlighted in black and demarcated by dashed lines. The *Q.Rg* values were < 1.3 and *Rg* was calculated as 86 Å and 167 Å, respectively. (**e**) SAXS Guinier analysis to determine the radius of gyration of the cross-section *(Rc);* linear fits are shown in red, with the fitted data range highlighted in black and demarcated by dashed lines. The *Q.Rc* values were < 1.3 and *Rc* was calculated as 16.6 Å and 15.3 Å, respectively.

## References

1. Hunter, N. Meiotic Recombination: The Essence of Heredity. Cold Spring Harb Perspect Biol 7(2015).

2. Zickler, D. & Kleckner, N. Recombination, Pairing, and Synapsis of Homologs during Meiosis. Cold Spring Harb Perspect Biol 7(2015).

3. Cahoon, C.K. & Hawley, R.S. Regulating the construction and demolition of the synaptonemal complex. Nat Struct Mol Biol 23, 369–77 (2016).

4. Kouznetsova, A., Benavente, R., Pastink, A. & Hoog, C. Meiosis in mice without a synaptonemal complex. PLoS One 6, e28255 (2011).

5. Sanchez-Saez, F. et al. Meiotic chromosome synapsis depends on multivalent SYCE1-SIX6OS1 interactions that are disrupted in cases of human infertility. Sci Adv 6(2020).

6. Geisinger, A. & Benavente, R. Mutations in Genes Coding for Synaptonemal Complex Proteins and Their Impact on Human Fertility. Cytogenet Genome Res 150, 77–85 (2016).

7. Baudat, F. & de Massy, B. Regulating double-stranded DNA break repair towards crossover or non-crossover during mammalian meiosis. Chromosome Res 15, 565–77 (2007).

8. MacGregor, I.A., Adams, I.R. & Gilbert, N. Large-scale chromatin organisation in interphase, mitosis and meiosis. Biochem J 476, 2141–2156 (2019).

9. Patel, L. et al. Dynamic reorganization of the genome shapes the recombination landscape in meiotic prophase. Nat Struct Mol Biol 26, 164–174 (2019).

10. Wang, Y. et al. Reprogramming of Meiotic Chromatin Architecture during Spermatogenesis. Mol Cell 73, 547–561 e6 (2019).

11. Schalbetter, S.A., Fudenberg, G., Baxter, J., Pollard, K.S. & Neale, M.J. Principles of meiotic chromosome assembly revealed in S. cerevisiae. Nat Commun 10, 4795 (2019).

12. Martini, E., Diaz, R.L., Hunter, N. & Keeney, S. Crossover homeostasis in yeast meiosis. Cell 126, 285–95 (2006).

13. Moses, M.J. Chromosomal structures in crayfish spermatocytes. J Biophys Biochem Cytol 2, 215–8 (1956).

14. Moses, M.J. Synaptinemal Complex. Annu Rev Genet 2, 363–412 (1968).

15. Westergaard, M. & von Wettstein, D. The synaptinemal complex. Annu Rev Genet 6, 71–110 (1972).

16. Solari, A.J. Synaptosomal complexes and associated structures in microspread human spermatocytes. Chromosoma 81, 315–37 (1980).

17. Spindler, M.C., Filbeck, S., Stigloher, C. & Benavente, R. Quantitative basis of meiotic chromosome synapsis analyzed by electron tomography. Sci Rep 9, 16102 (2019).

18. Solari, A.J. & Moses, M.J. The structure of the central region in the synaptonemal complexes of hamster and cricket spermatocytes. J Cell Biol 56, 145–52 (1973).

19. Schmekel, K., Skoglund, U. & Daneholt, B. The three-dimensional structure of the central region in a synaptonemal complex: a comparison between rat and two insect species, Drosophila melanogaster and Blaps cribrosa. Chromosoma 102, 682–92 (1993).

20. Schucker, K., Holm, T., Franke, C., Sauer, M. & Benavente, R. Elucidation of synaptonemal complex organization by super-resolution imaging with isotropic resolution. Proc Natl Acad Sci U S A 112, 2029–33 (2015).

21. Schmekel, K. et al. Organization of SCP1 protein molecules within synaptonemal complexes of the rat. Exp Cell Res 226, 20–30 (1996).

22. Liu, J.G. et al. Localization of the N-terminus of SCP1 to the central element of the synaptonemal complex and evidence for direct interactions between the N-termini of SCP1 molecules organized head-to-head. Exp Cell Res 226, 11–9 (1996).

23. Dunce, J.M. et al. Structural basis of meiotic chromosome synapsis through SYCP1 self-assembly. Nat Struct Mol Biol 25, 557–569 (2018).

24. Yuan, L. et al. Female germ cell aneuploidy and embryo death in mice lacking the meiosis-specific protein SCP3. Science 296, 1115–8 (2002).

25. Yuan, L. et al. The murine SCP3 gene is required for synaptonemal complex assembly, chromosome synapsis, and male fertility. Mol Cell 5, 73–83 (2000).

26. Yang, F. et al. Mouse SYCP2 is required for synaptonemal complex assembly and chromosomal synapsis during male meiosis. J Cell Biol 173, 497–507 (2006).

27. Costa, Y. et al. Two novel proteins recruited by synaptonemal complex protein 1 (SYCP1) are at the centre of meiosis. J Cell Sci 118, 2755–62 (2005).

28. Schramm, S. et al. A novel mouse synaptonemal complex protein is essential for loading of central element proteins, recombination, and fertility. PLoS Genet 7, e1002088 (2011).

29. Hamer, G. et al. Characterization of a novel meiosis-specific protein within the central element of the synaptonemal complex. Journal of Cell Science 119, 4025–4032 (2006).

30. Gomez, H.L. et al. C14ORF39/SIX6OS1 is a constituent of the synaptonemal complex and is essential for mouse fertility. Nat Commun 7, 13298 (2016).

31. de Vries, F.A. et al. Mouse Sycp1 functions in synaptonemal complex assembly, meiotic recombination, and XY body formation. Genes Dev 19, 1376–89 (2005).

32. Bolcun-Filas, E. et al. Mutation of the mouse Syce1 gene disrupts synapsis and suggests a link between synaptonemal complex structural components and DNA repair. PLoS Genet 5, e1000393 (2009).

33. Bolcun-Filas, E. et al. SYCE2 is required for synaptonemal complex assembly, double strand break repair, and homologous recombination. Journal of Cell Biology 176, 741–747 (2007).

34. Hamer, G. et al. Progression of meiotic recombination requires structural maturation of the central element of the synaptonemal complex. J Cell Sci 121, 2445–51 (2008).

35. Fraune, J., Schramm, S., Alsheimer, M. & Benavente, R. The mammalian synaptonemal complex: Protein components, assembly and role in meiotic recombination. Ėxp Cell Res (2012).

36. Lu, J. et al. Structural insight into the central element assembly of the synaptonemal complex. Sci Rep 4, 7059 (2014).

37. Davies, O.R., Maman, J.D. & Pellegrini, L. Structural analysis of the human SYCE2-TEX12 complex provides molecular insights into synaptonemal complex assembly. Open Biology 2, 120099 (2012).

38. Syrjanen, J.L., Pellegrini, L. & Davies, O.R. A molecular model for the role of SYCP3 in meiotic chromosome organisation. Ėlife 3(2014).

39. Syrjanen, J.L. et al. Single-molecule observation of DNA compaction by meiotic protein SYCP3. Ėlife 6(2017).

40. Dunne, O.M. & Davies, O.R. A molecular model for self-assembly of the synaptonemal complex protein SYCE3. J Biol Chem 294, 9260–9275 (2019).

41. Dunne, O.M. & Davies, O.R. Molecular structure of human synaptonemal complex protein SYCE1. Chromosoma (2019).

42. West, A.M. et al. A conserved filamentous assembly underlies the structure of the meiotic chromosome axis. Ėlife 8(2019).

43. Bollschweiler, D. et al. Molecular architecture of the SYCP3 fibre and its interaction with DNA. Open Biol 9, 190094 (2019).

44. Yuan, L. et al. The synaptonemal complex protein SCP3 can form multistranded, cross-striated fibers in vivo. J Cell Biol 142, 331–9 (1998).

45. Baier, A., Alsheimer, M. & Benavente, R. Synaptonemal complex protein SYCP3: Conserved polymerization properties among vertebrates. Biochim Biophys Acta 1774, 595–602 (2007).

46. Baier, A., Alsheimer, M., Volff, J.N. & Benavente, R. Synaptonemal complex protein SYCP3 of the rat: evolutionarily conserved domains and the assembly of higher order structures. Sex Dev 1, 161–8 (2007).

47. Ortiz, R. et al. Cytochemical study of the distribution of RNA and DNA in the synaptonemal complex of guinea-pig and rat spermatocytes. Eur J Histochem 46, 133–42 (2002).

48. Caballero, I. et al. ARCIMBOLDO on coiled coils. Acta Crystallogr D Struct Biol 74, 194–204 (2018).

49. Squire, J. Protein Conformation and Characterization. in The Structural Basis of Muscular Contraction 129–155 (Springer US, Boston, MA, 1981).

50. Kreplak, L., Doucet, J. & Briki, F. Unraveling double stranded alpha-helical coiled coils: an x-ray diffraction study on hard alpha-keratin fibers. Biopolymers 58, 526–33 (2001).

51. Er Rafik, M., Doucet, J. & Briki, F. The intermediate filament architecture as determined by X-ray diffraction modeling of hard alpha-keratin. Biophys J 86, 3893–904 (2004).

52. Busson, B., Briki, F. & Doucet, J. Side-chains configurations in coiled coils revealed by the 5.15-A meridional reflection on hard alpha-keratin X-ray diffraction patterns. J Struct Biol 125, 1–10 (1999).

53. Bai, Y., Luo, Q. & Liu, J. Protein self-assembly via supramolecular strategies. Chem Soc Rev 45, 2756–67 (2016).

54. Garcia-Seisdedos, H., Empereur-Mot, C., Elad, N. & Levy, E.D. Proteins evolve on the edge of supramolecular self-assembly. Nature 548, 244–247 (2017).

55. McManus, J.J., Charbonneau, P., Zaccarelli, E. & Asherie, N. The physics of protein self-assembly. Current Opinion in Colloid & Interface Science 22, 73–79 (2016).

56. Ahn, J. et al. Structural basis for lamin assembly at the molecular level. Nat Commun 10, 3757 (2019).

57. Aziz, A. et al. The structure of vimentin linker 1 and rod 1B domains characterized by site-directed spin-labeling electron paramagnetic resonance (SDSL-EPR) and X-ray crystallography. J Biol Chem 287, 28349–61 (2012).

58. Pang, A.H., Obiero, J.M., Kulczyk, A.W., Sviripa, V.M. & Tsodikov, O.V. A crystal structure of coil 1B of vimentin in the filamentous form provides a model of a high-order assembly of a vimentin filament. FEBS J 285, 2888–2899 (2018).

59. Chernyatina, A.A., Nicolet, S., Aebi, U., Herrmann, H. & Strelkov, S.V. Atomic structure of the vimentin central alpha-helical domain and its implications for intermediate filament assembly. Proc Natl Acad Sci U S A 109, 13620–5 (2012).

60. Lee, C.H., Kim, M.S., Chung, B.M., Leahy, D.J. & Coulombe, P.A. Structural basis for heteromeric assembly and perinuclear organization of keratin filaments. Nat Struct Mol Biol 19, 707–15 (2012).

61. Bunick, C.G. & Milstone, L.M. The X-Ray Crystal Structure of the Keratin 1-Keratin 10 Helix 2B Heterodimer Reveals Molecular Surface Properties and Biochemical Insights into Human Skin Disease. J Invest Dermatol 137, 142–150 (2017).

62. Eldirany, S.A., Ho, M., Hinbest, A.J., Lomakin, I.B. & Bunick, C.G. Human keratin 1/10-1B tetramer structures reveal a knob-pocket mechanism in intermediate filament assembly. EMBO J 38(2019).

63. Eldirany, S.A., Lomakin, I.B., Ho, M. & Bunick, C.G. Recent insight into intermediate filament structure. Curr Opin Cell Biol 68, 132–143 (2020).

64. Koster, S., Weitz, D.A., Goldman, R.D., Aebi, U. & Herrmann, H. Intermediate filament mechanics in vitro and in the cell: from coiled coils to filaments, fibers and networks. Curr Opin Cell Biol 32, 82–91 (2015).

65. Herrmann, H. & Aebi, U. Intermediate Filaments: Structure and Assembly. Cold Spring Harb Perspect Biol 8(2016).

66. Turgay, Y. et al. The molecular architecture of lamins in somatic cells. Nature 543, 261–264 (2017).

67. Kabsch, W. Xds. Acta Crystallogr D Biol Crystallogr 66, 125–32 (2010).

68. Diederichs, K., McSweeney, S. & Ravelli, R.B. Zero-dose extrapolation as part of macromolecular synchrotron data reduction. Acta Crystallogr D Biol Crystallogr 59, 903–9 (2003).

69. Tickle, I.J. et al. STARANISO (http://staraniso.globalphasing.org/cgi-bin/staraniso.cgi). Cambridge, United Kingdom: Global Phasing Ltd. (2018).

70. Rodriguez, D.D. et al. Crystallographic ab initio protein structure solution below atomic resolution. Nature Methods 6, 651–U39 (2009).

71. McCoy, A.J. et al. Phaser crystallographic software. Journal of Applied Crystallography 40, 658–674 (2007).

72. Adams, P.D. et al. PHENIX: a comprehensive Python-based system for macromolecular structure solution. Acta Crystallogr D Biol Crystallogr 66, 213–21 (2010).

73. Emsley, P., Lohkamp, B., Scott, W.G. & Cowtan, K. Features and development of Coot. Acta Crystallogr D Biol Crystallogr 66, 486–501 (2010).

74. Chen, V.B. et al. MolProbity: all-atom structure validation for macromolecular crystallography. Acta Crystallographica Section D-Biological Crystallography 66, 12–21 (2010).

75. Vonrhein, C. et al. Data processing and analysis with the autoPROC toolbox. Acta Crystallogr D Biol Crystallogr 67, 293–302 (2011).

76. Sreerama, N. & Woody, R.W. Estimation of protein secondary structure from circular dichroism spectra: comparison of CONTIN, SELCON, and CDSSTR methods with an expanded reference set. Anal Biochem 287, 252–60 (2000).

77. Whitmore, L. & Wallace, B.A. Protein secondary structure analyses from circular dichroism spectroscopy: methods and reference databases. Biopolymers 89, 392–400 (2008).

78. Schindelin, J. et al. Fiji: an open-source platform for biological-image analysis. Nat Methods 9, 676–82 (2012).

79. P.V. Konarev, V.V.V., A.V.Sokolova, M.H.J. Koch, D. I. Svergun. PRIMUS – a Windows-PC based system for small-angle scattering data analysis. J Appl Cryst. 36, 1277–1282 (2003).

80. Franke, D. & Svergun, D.I. DAMMIF, a program for rapid ab-initio shape determination in small-angle scattering. J Appl Crystallogr 42, 342–346 (2009).

81. Svergun, D.I. Restoring low resolution structure of biological macromolecules from solution scattering using simulated annealing. Biophys J 76, 2879–86 (1999).

82. Kozin, M.B. & Svergun, D.I. Automated matching of high- and low-resolution structural models. Journal of Applied Crystallography 34, 33–41 (2001).

83. Svergun D.I., B.C.K.M.H.J. CRYSOL – a Program to Evaluate X-ray Solution Scattering of Biological Macromolecules from Atomic Coordinates. J. Appl. Cryst. 28, 768–773. (1995).

84. Schneidman-Duhovny, D., Hammel, M., Tainer, J.A. & Sali, A. FoXS, FoXSDock and MultiFoXS: Single-state and multi-state structural modeling of proteins and their complexes based on SAXS profiles. Nucleic Acids Res 44, W424–9 (2016).

85. Petoukhov, M.V. et al. New developments in the ATSAS program package for small-angle scattering data analysis. J Appl Crystallogr 45, 342–350 (2012).

86. Wood, C.W. & Woolfson, D.N. CCBuilder 2.0: Powerful and accessible coiled-coil modeling. Protein Sci 27, 103–111 (2018).

87. Nivon, L.G., Moretti, R. & Baker, D. A Pareto-optimal refinement method for protein design scaffolds. PLoS One 8, e59004 (2013).

88. Waterhouse, A.M., Procter, J.B., Martin, D.M., Clamp, M. & Barton, G.J. Jalview Version 2--a multiple sequence alignment editor and analysis workbench. Bioinformatics 25, 1189–91 (2009).

